# Epstein-Barr virus induced epigenetic reprogramming drives cancer stem cell emergence in breast cancer

**DOI:** 10.64898/2026.03.10.710870

**Authors:** Bernard Friedenson

**Affiliations:** Department of Biochemistry and Molecular Genetics and Illinois Cancer Center, College of Medicine University of Illinois Chicago Chicago, Illinois

## Abstract

Basal breast cancers display stem cell associated basal programs, but originate from luminal progenitors. This lineage paradox may be created by Epstein–Barr virus (EBV) infection. Across more than 2,000 breast cancer genomes, coordinated methyla-tion changes appeared in cis-regulatory elements governing stem cell differentiation. Methylation positions followed EBV-associated malignancies with striking accuracy independent of whether ER-status marked a luminal or basal cancer. EBV-driven epigenetic reprogramming was incompatible with tumor infiltrating lymphocytes and disrupted lineage specification before tumorigenesis. Breast cancers commonly showed coordinated viral response indicators that tracked with antigen presentation and stem cell differentiation programs. Non-malignant keratinocytes with resolved EBV infections retained some aberrantly methylated loci. Analyses of non-EBV skin carcinoma, randomized genomic sites, endogenous retroelements, DUX4, and repli-cation clocks confirmed the specificity of EBV-linked alterations. These findings posi-tion EBV as a developmental lineage hijacker that reprograms cells into premalignant stem-like states.

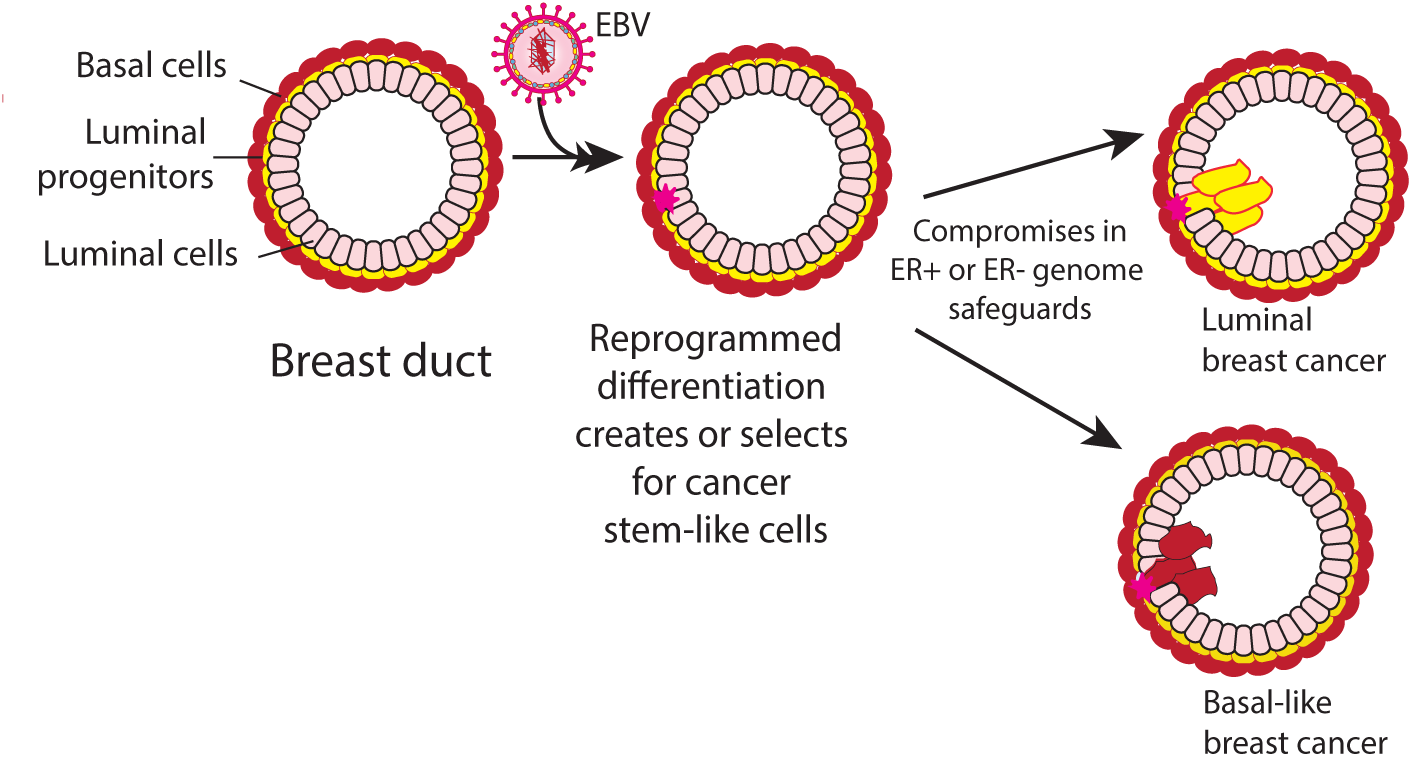

EBV infection reprograms the methylome of breast luminal progenitors to generate or select for cancer stem-like cells.

**In Brief:** Epstein-Barr virus (EBV) rewires developmental programs that specify cell identity, creating cancer stem-like cells with malignant potential. This targeted reshaping of progenitors links viral infection to the earliest steps of breast cancer development.

**Highlights:**

1. Epstein-Barr virus (EBV) infection reprograms the methylome of lu-minal progenitors generating breast cancer stem-like cells.
2. EBV-driven methylation sites overlap luminal and basal breast cancer signatures that regulate stem cell differentiation.
3. Characteristic responses to viral infection are common in breast can-cer and they correlate with gene programs needed to differentiate progenitor cells.
4. Even after viral clearance, EBV leaves persistent methylation scars in non-malignant cells that may increase risks for future malignancy.

## Introduction

Basal-like breast cancers are among the most aggressive tumor subtypes, marked by stem-like cellular features and poor clinical outcomes. Yet a single-cell transcriptomic atlas of over 700,000 normal breast cells identified no canonical stem-cell population in the basal layer ^1^, even though small malignant cell foci expressing basal cell markers appear in luminal and HER2-enriched cancers ^2–4^. This paradox raises a fundamental question: if the normal basal layer lacks a discrete stem-cell pool, how do stem-like states emerge in tumors?

Loss of BRCA1 function, epigenomic disruptions ^5^, and oncogenic hits accompany luminal progenitors progressively acquiring basal programs ^6^, but whether there is a central underlying cause has not been studied. One possibility is epigenetic reprogramming by an upstream viral pathogen. Epstein-Barr virus (EBV) can infect mammary epithelial cells and promote malignant traits in vitro ^7^, but the physiological relevance of this finding is uncertain. The experiments relied on engineered culture systems lacking the layered architecture of breast ducts; examined only a limited gene set; and did not examine epigenetic consequences. Infection occurred mainly in myoepithelial cells, required CD21-receptors that are rare in mammary epithelium, and ER-positive cell lines were resistant. Moreover, a widely held opinion is that EBV primarily infects an occasional tumor infiltrating lymphocyte (TIL), not the tumor itself ^8^. These limitations underscore the need for an EBV focused genome and epigenome wide analyses in primary human tumors. Such studies are essential to determine whether viral reprogramming contributes to cancer stem-like phenotypes.

Both breast cancers ^9^ and EBV-associated cancers ^10^ exhibit cancer stem cell hierarchies. A unifying model is that these hierarchies arise because EBV disrupts normal stem cell lineages, generating or selecting for basal-like stem cells in contexts where they do not normally exist. Under this model, EBV gains a selective advantage when infected stem cells cannot be restored to their uninfected states. Infection would therefore select for stem cells with impaired differentiation or self renewal. If correct, molecular lesions that perturb stem cell lineages in breast cancer should resemble those seen in EBV-associated cancers.

Multiple lines of evidence support such a connection. EBV disrupts epithelial homeostasis, induces genomic instability, and extensively manipulates host methylation during latency and transformation. Viral genomes acquire nucleosomes tightly coupled to methylation as early as two days post infection ^11^. EBV encoded transcription factors profoundly alter host gene expression, accompanied by large scale chromatin restructuring^11^. In breast cancer systems, strong correlations have been established between methylation sites and nearby genes (cis-methylation) ^12^. These correlating methylation patterns permit testing their effects on breast cancer stem cell hierarchies. Adult stem cell replication both shapes and responds to methylation instability ^13^.

If EBV can reprogram breast luminal progenitor cells, then breast tumors should retain methylation patterns resembling those in EBV-associated cancers—especially at regulatory loci that control early lineage commitment decisions. One way to estimate how these effects play out in different breast cell types is to compare ER-positive (luminal) and ER-negative (basal) tumors, which serve as rough proxies for luminal derived versus basal enriched disease. EBV-like methylation in these stem cell pools could leave durable genomic and epigenomic scars, predisposing cells to malignant progression even after the viral genome is no longer present.

In known EBV-driven cancers, the virus reprograms stem cell pathways required for tissue maintenance and regeneration as it induces massive host cell methylation changes and reshapes chromatin accessibility ^14,15^. Methylation patterns are strong predictors of such reprogramming ^16,17^. In the present study, methylation profiles from over 2,000 breast cancer genomes revealed alterations in cis-regulatory elements governing stem cell differentiation, function, and microenvironmental interactions. These methylation sites were absent in normal breast tissue but appeared in identical or virtually identical positions in prototypical EBV-associated cancers. Striking agreement between breast cancers and EBV cancers occurred in completely unrelated data, produced in different laboratories sometimes decades apart. The correlation was so strong and so consistent with established EBV biology that it supports the hypothesis that EBV selects for epigenetic modifications in stem cell lineages.

Breast cancer stem cell related methylation loci shared with EBV cancers include genes, morphogens, lineage specification factors, and regulators of luminal tight junctions. Antiviral immune response pathways were especially targeted, showing coordinated effects in hallmark antiviral response genes and relationships to differentiation pathways.

Together these findings recast EBV not merely as an oncogenic virus, but as a developmental disruptor that hijacks epigenetic machinery to create and select for premalignant stem-like states. Because EBV infection is virtually universal and often acquired early in life, these selected signatures likely precede tumorigenesis. Persistent methylation abnormalities in non-malignant keratinocytes after resolved EBV infection further suggest durable epigenetic memory and potential long-term risk. Analyses of tumor infiltrating lymphocytes, non-EBV skin carcinoma, randomized genomic sites, endogenous retroelements, *DUX4* genes and replication clocks confirmed the specificity of EBV-linked alterations.

## Results

### Abnormal methylation sites in breast cancer that regulate key genes align closely with those in EBV-associated cancers

For all 22 human autosomes, minimum distances were calculated between abnormal methylation at breast cancer gene control elements and the EBV-associated cancers nasopharyngeal cancer (NPC) and Burkitt lymphoma (BL). The genome coordinates for methyl groups shared between breast cancer and EBV cancers were sometimes exactly the same, but the most frequent agreement was within 10 bp (Fig. 1). For example, on chromosome 2, fifteen of 2,319 NPC methylation sites were within 10 bp of the 789 breast cancer sites, far exceeding the 0.158 matches expected by chance (Poisson p ≈ 1.3 × 10^-^²⁷).

**Fig. 1.**
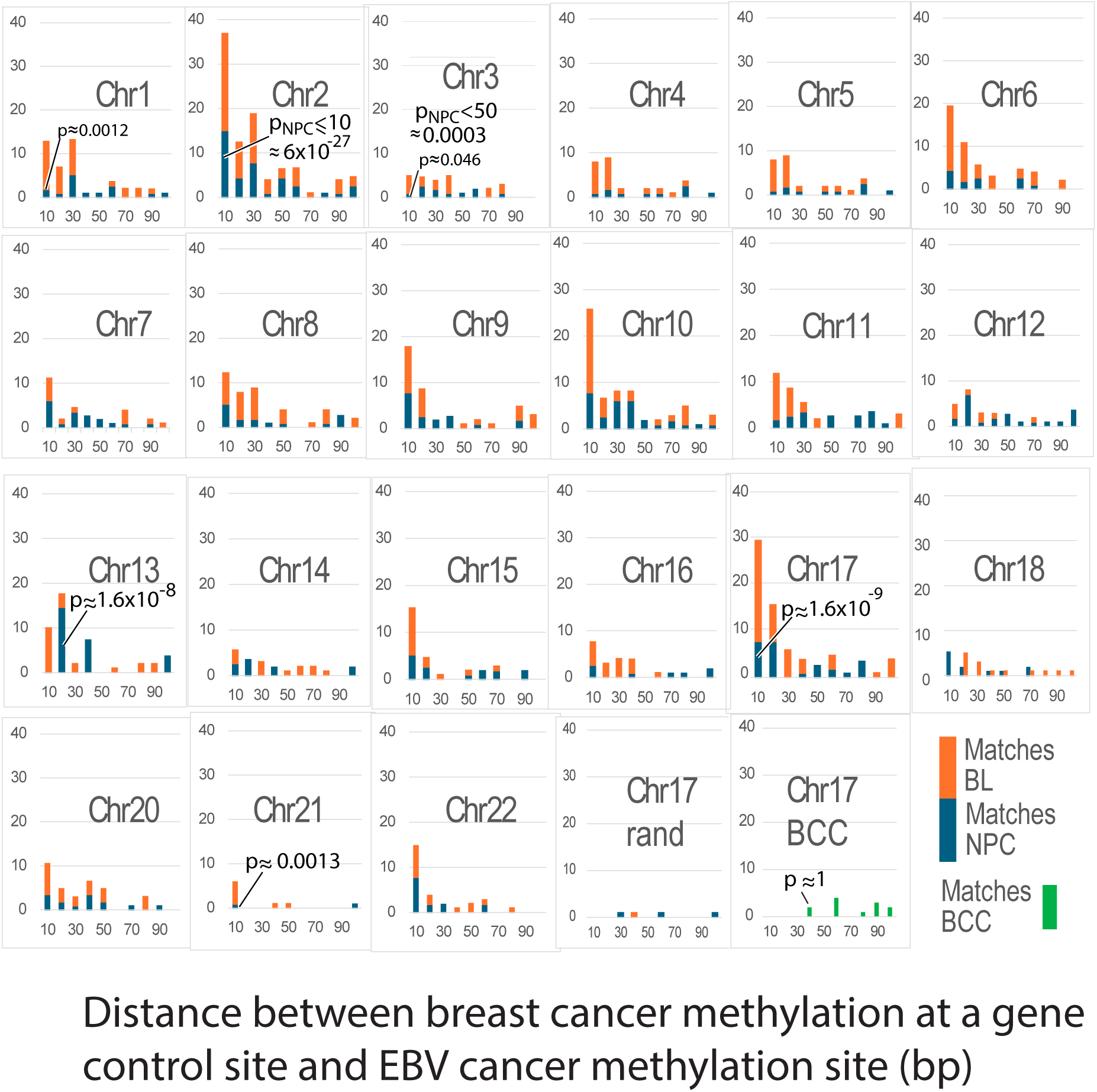
Minimum distance between abnormal methylation sites in breast cancer and EBV cancers. Across most autosomes, abnormal methylation sites in breast cancer lie within 10 bp of abnormal sites in the EBV driven cancers NPC and BL. For visualization, NPC and BL data were normalized to the same total numbers of patients; however, statistical tests for NPC were performed using the orig-inal, unnormalized NPC totals. A control comparison on chromosome 17 shows that breast cancer sites rarely match a random distribution. One representative trial (Chr17 rand) produced almost no matches, and other trials were blank. The non-EBV cancer (skin) BCC (green bars) also shows a pattern on chromosome 17 that does not resemble the close agreement observed between breast cancer and the EBV-associated cancers.

Chromosome 17 showed similar enrichment, with nine matches within 10 bp (p ≈ 2 × 10^-^⁷). Even on chromosome 3, where matching methylated sites were sparse, five NPC–breast cancer sites agreed within 50 bp (p ≈ 0.0003).

Distribution level comparisons also supported concordance. Mann–Whitney tests on chromosomes 2, 7, 9-12, and 20 indicated no statistically significant differences in the meth-ylation distributions between breast cancer and NPC (p>0.05). For chromosomes 9–11, the confidence intervals for the median differences included zero, which further supports their similarity. Even for chromosomes with only a few close matches, the observed distributions were statistically indistinguishable.

Control analyses confirmed specificity. Randomized datasets produced blank or nearly blank patterns (Fig. 1, lower right). A second control compared 13,766 CpGs on chromosome 17 from (skin) basal cell carcinoma (BCC)^18^ to breast cancer methylation positions. Far fewer and weaker positional matches were found. Mann–Whitney testing confirmed that BCC and breast cancer distributions differed significantly (p<0.0001).

Together, these results show that breast cancer chromosomes frequently share abnormal methylation positions with EBV-driven cancers. Shared methylation at such specific gene regulatory sites may be crucial for producing both cancer types.

### Breast cancer methylation loci that match EBV-cancers represent selection for stem cell genes

To test whether shared methylation sites preferentially targeted specific biological functions, cis-methylation controlled genes in breast cancers^12^ were compared to loci also methylated in EBV cancers. Chromosome 17 was examined first because it encodes *BRCA1* and *ERBB2* (*HER2*) genes, both significant in breast cancer. In the breast cancer cohort, chromosome 17 had 405 gene entries with abnormally methylated cis-regulatory sites with 85% <=200 bp long. As shown in Fig. 2, comparisons of abnormal methylation sites on chromosome 17 in breast cancer ^12^, NPC ^14^, and BL ^19^ converged on the same functions.

**Fig. 2.**
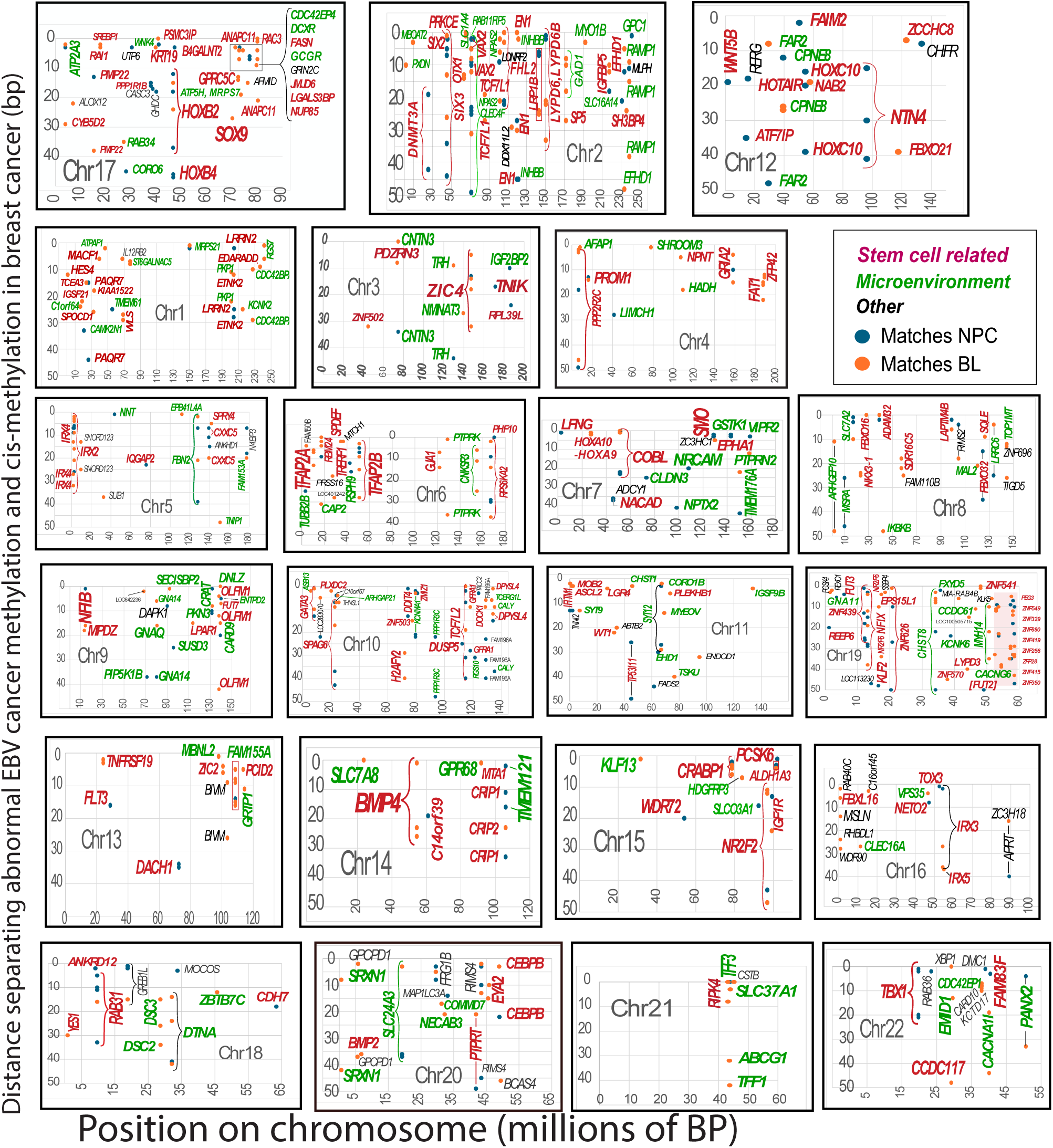
Methylation patterns in regulatory regions of breast cancer stem cell genes closely parallel those in the EBV associated cancers NPC and BL. Blue and orange dots (see second row legend) mark abnormal methylation sites in NPC and BL that fall within 50 bp of abnormal cis methylation sites in breast cancer. Genes with stem cell–related functions are shown in red, genes influencing the microenvironment in green, and genes with other functions in black. Many breast cancer regulatory sites align at identical or near identical positions in EBV cancers, producing the dense clusters at the top of each panel. The three chromosomes in the top row are discussed in the text. On chromosome 19, the red shaded region highlights zinc finger gene matches, although the underlying data for this cluster are incomplete.

For instance, the three different cancers each targeted regulatory regions of *SOX9. SOX9* counterbalances the core stem-cell gene *SOX2*, which governs pluripotency. Together *SOX9* and *SOX2* shape cancer cell plasticity and metastasis ^20^. *SOX9* also participates in the *NOTCH* signaling network governing mammary stem cell self-renewal and lineage choice ^21^. *KRT19,* a *NOTCH* signaling control point, was methylated within three bases of the same position in breast cancer and BL. Breast cancer and EBV-associated tumors shared hypermethylated clusters within the *HOXB* locus, including regions near *HOXB2* and *HOXB4*. These blocks overlapped regulatory elements characteristic of stem cell–related chromatin states ^22^.

*HOXB* genes have documented roles in both breast and EBV-associated cancers^23,24^.

Finding *SOX9, KRT19,* and *HOX* genes with nearly identical abnormal methylation positions in breast and EBV-driven cancers suggested that EBV-induced methylation had disrupted breast stem cell differentiation. Fig. 1 hints at this possibility: children with BL consistently exhibited more abnormal EBV-matching methylation than adults with NPC consistent with the age related decline in stem cell numbers and lineage potential. This pattern prompted broader analyses on additional breast cancer autosomes.

Abnormal methylation positions shared between breast cancer and EBV-associated cancers extended throughout the genome with a strong bias toward stem-cell related genes and lineage defining transcription factors. Supplementary Table S1 summarizes the functions of ∼350 of these genes. As highlighted in Fig. 2, many breast cancer gene promoters and enhancer elements reside within 50 bp of EBV-cancer methylation sites, disrupting morphogen responsive *HOX* genes and stem cell linked regulators.

On chromosome 17 (Figs. 2, 3; Table S1), ∼66% of abnormally methylated cis-regulatory positions in breast cancer coincide with EBV cancer methylation loci. Across all autosomes, 48% of coinciding genes participate in stem cell programs or differentiation pathways; representing widespread methylation changes in regulators of cell-type commitment, self renewal, and tissue homeostasis.

For example, on chromosome 2, genes with altered controls in both breast and EBV-cancers, (including *HOX, SIX2, SIX3, VAX2,* and *EN1*), have strong developmental and stem cell connections (Figs 2, 3; Table S1). *HOX* genes set broad positional identity ^25^, *SIX, VAX,* and *EN* genes refine regional patterning and morphogen gradient responses essential for tissue homeostasis ^26^. Chromosome 12 results include an RNA-based mechanism. The long noncoding RNA *HOTAIR*, a 2.2 kb antisense transcript, represses *HOXD* cluster transcription by recruiting chromatin modifying complexes ^27^.

Table S1 documents 167 instances of abnormal methylation in stem cell related pathways in breast cancer that match patterns seen in NPC or BL within 50 bp. Collectively, these findings suggest that EBV-driven reprogramming of stem-cell pathways is a shared feature across all three cancers.

### Shared abnormal methylation loci regulate multiple cell differentiation pathways

To assess EBV-linked methylation disruption of stem cell lineage, differentiation pathways with EBV-related methylated gene controls were assessed. Six chromosomes were examined that encoded *HOX* gene clusters and had numerous close (<10 bp) matches to EBV cancer methylation sites. As shown in Fig. 3, morphogen associated pathways were recurrently impacted, with specific pathway sets differing by chromosome. On chromo-some 6, *OCT4* and *SOX2* were the most frequent targets. These core stem cell regulators act cooperatively to reprogram somatic cells and maintain pluripotency. Their deregulation can collapse the entire pluripotency network ^28,29^. *HOX* regulatory elements on four chro-mosomes were methylated. Additional genes connected to *Wnt* and other stem cell differ-entiation pathways are also frequent targets. Together, these findings indicate that EBV-re-lated methylation patterns can substantially reprogram stem cell differentiation, potentially compromising tissue regeneration.

**Fig. 3.**
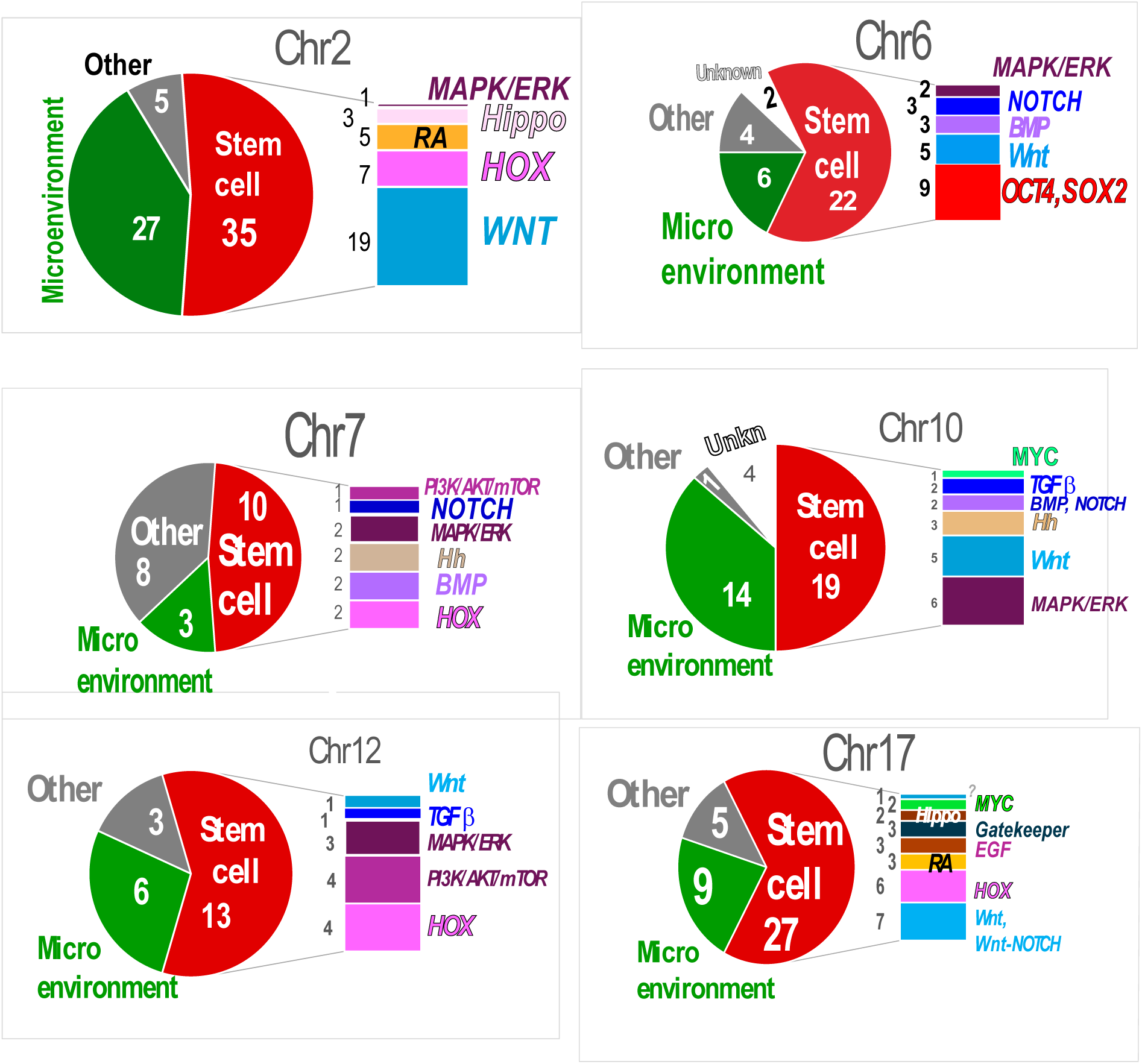
Breast cancer gene control regions with abnormal cis-methylation corresponding to EBV cancers affect stem cell differentiation pathways and morphogenesis. Pie charts for data from mul-tiple chromosomes indicate the proportion of shared abnormal methylation events affecting stem cell programs, the microenvironment, or other functions. Bar charts to the right show the differentiation pathways most frequently impacted by breast cancer methylation sites that coincide with EBV can-cers; the counts for genes representing each pathway appear on the left of each bar. On chromosome 6, shared abnormal methylation directly targets the core pluripotency regulators *OCT4* and *SOX2*. Most methylated sites mapped to multiple pathways, but only the predominant morphogen connected path-ways are highlighted.

### Shared methylation loci dysregulate expression of stem cell genes and stem cell markers

In breast cancer, cis-specific methylation and gene expression correlate “at hundreds of promoters and over a thousand distal elements” ^12^ often at large genome distances.

Fig. 4A presents, for each autosome, the distribution of 356 genes located adjacent to breast cancer methylation sites that coincide within 50 bp of corresponding abnormal methylation sites in EBV-driven cancers. Expression data ^30^ were available for 345 of the 356 genes and all 345 exhibited significant abnormalities across multiple breast cancers (Fig. 4B).

**Fig. 4A.**
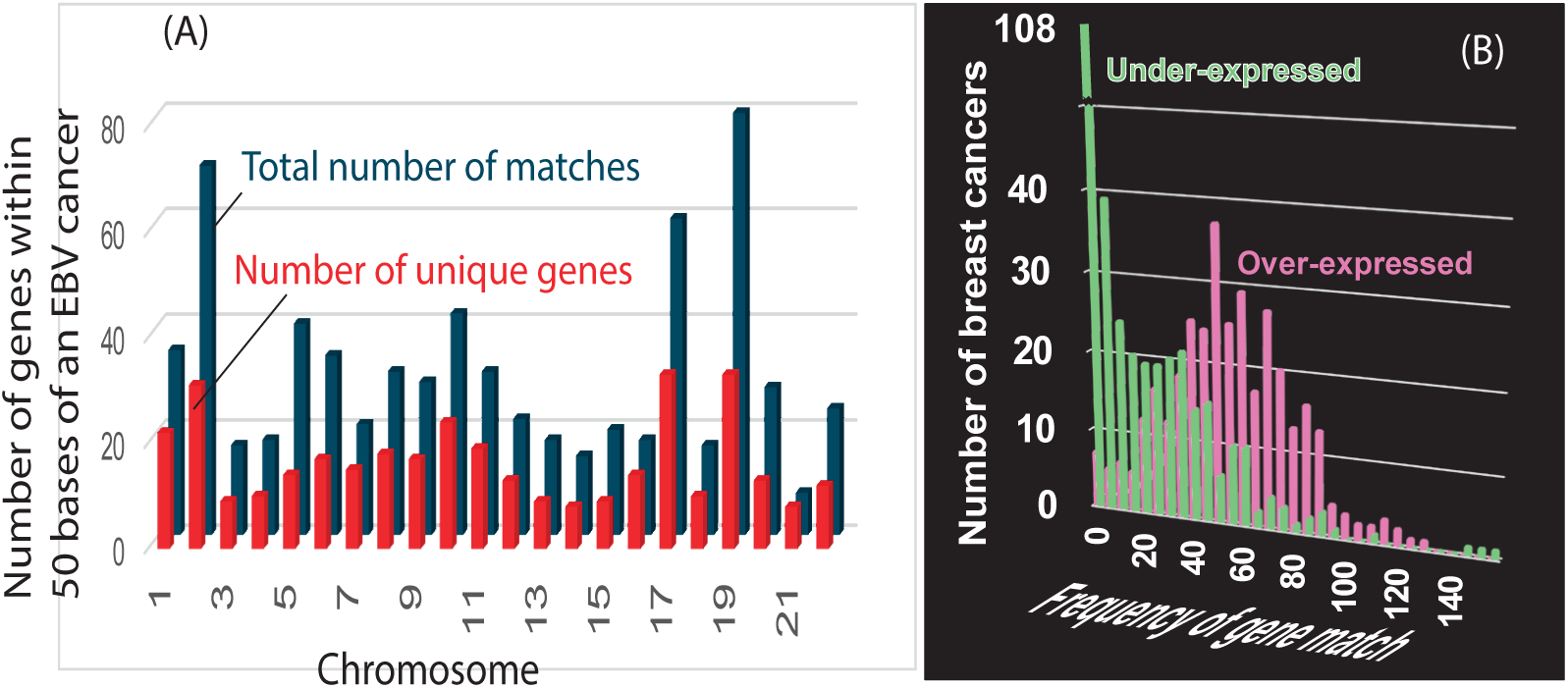
Variation in number of genes controlled at sites that match (<50 bp) EBV-associated cancer methylation sites. Unique genes are in red. Multiple matches to the same gene regulatory sites are in blue. **B. Genes controlled by methylation at sites that match EBV cancers are likely expressed at abnormal levels.**

**Fig. 4C.**
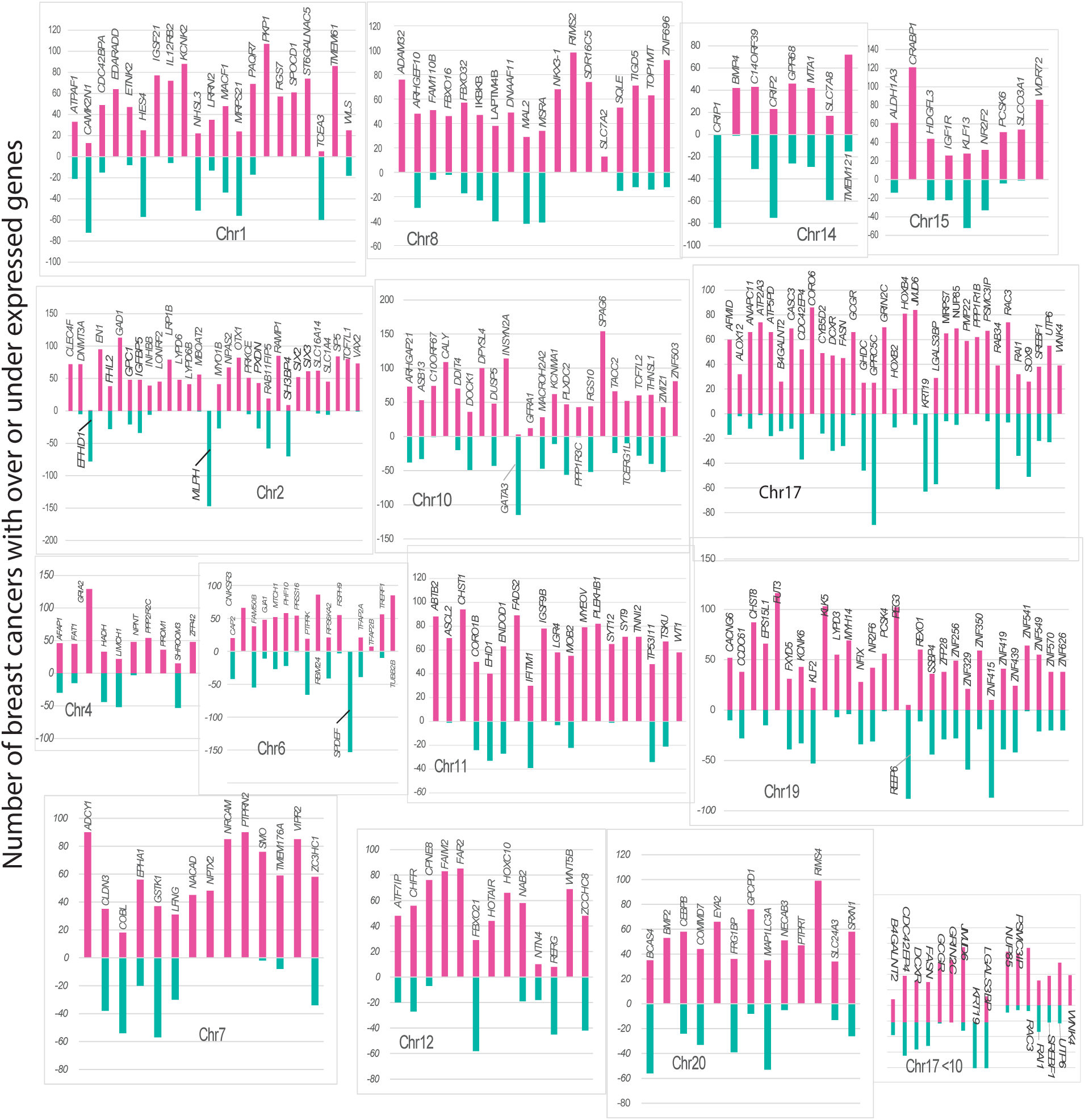
Genes with abnormal methylation in breast cancers that match EBV-associated cancers within <50 bp show significant over-or under-expression across multiple autosomes (|Z-score| > 2). The vertical axis indicates how many breast cancers in the cohort exhibit abnormal methylation for each indicated gene. Many of the over-expressed genes are not expressed in immune cells, making explanations based on TILs unlikely. Within each chromosome panel, genes are listed alphabetically. Many regulatory regions sit <10 bp from an EBV-methylated site, as illustrated for chromosome 17 (lower right). Red bars denote over-expressed genes; green bars denote under-expressed genes.

Fig. 4C summarizes the numbers of tumors with dysregulated expression of breast cancer genes near EBV-methylation sites, and Table 1 highlights representative genes grouped by their roles in stem cell related differentiation pathways. Dysregulation of these genes influences hundreds of downstream targets governing stem cell differentiation and cancer stem cell states ^31^.

**Table 1.** Effects on differentiation by over-and under-expressed genes in breast cancers. Abnormal methylation of these gene control sites closely relates to abnormal methylation sites in EBV cancers

| Gene | Chr | General function | Expression in breast cancers | Context and reference |
| --- | --- | --- | --- | --- |
| **BMP2** | 20 | Morphogen | Over-expressed | Drives differentiation |
| **BMP4** | 14 | Morphogen | Over-expressed | Drives differentiation |
| **CRABP1** | 15 | Morphogen | Over-expressed | Powerful regulator of stem cell identity specification via retinoic acid signaling, diminishes neural and embryonic stem cell expansion <sup>70</sup>. |
| **GRIA2** | 4 | Differentiation regulator | Over-expressed | Neurodevelopment gene. Steers progenitors toward specific cell fates after an injury <sup>71</sup> |
| **HOTAIR** | 12 | Homeobox gene regulator | Over-expressed | Epigenetic regulator. causes H3K27 methylation; increases breast cancer invasiveness |
| **HOXC10** | 12 | Homeobox gene | Over-expressed | Activates *ERK*, *AKT*, *p65*, and EMT-related genes |
| **KLF2** | 19 | Pluripotency regulator <sup>72</sup>. | Over-expressed<br>Under-expressed | Increases expression of stem cell markers like OCT4 and NANOG, suppresses differentiation markers to maintain stemness and self-renewal <sup>73</sup> |
| **SIX2** | 2 | Homeobox gene | Over-expressed | Enhances expression of stem cell markers *SOX2* and *Nanog* <sup>74,75</sup> Activates Wnt/β-catenin and MAPK pathways; |
| **SIX3** | 2 | Homeobox gene | Over-expressed | Modulates Wnt signaling; enhances proliferation and migration in gastric cancer <sup>76</sup> |
| **SMO** | 7 | Morphogen-responsive gene | Over-expressed | Guides differentiation into specific identities |
| **SOX9** | 17 | Pluripotency regulator | Over-expressed<br>Under-expressed | Antagonizes and opposes pluripotency factor *SOX2* <sup>77</sup> |
| **SPAG6** | 10 | Differentiation regulator | Over-expressed | A cilia-related gene; over-expression biases stem cells toward specific lineages and inhibits proliferation of neural progenitors <sup>78</sup>. |
| **TCF7L1** | 2 | Pluripotency regulator vs. self renewal | Over-expressed | Silences Wnt targets; maintains pluripotent state preventing differentiation <sup>36</sup> |
| **TFAP2A** | 6 | Pluripotency exit regulator | Under-expressed<br>Over-expressed | Methylation locks cells in self-renewal; disrupts placental lineage commitment <sup>79</sup>. |
| ***TFAP2B*** | 6 | Differentiation regulator | Over-expressed (occasional) | Determines differentiation outcome in an embryonic melanocyte stem cell <sup>80</sup>. Suppresses Wnt/β-catenin pathways in certain cancers <sup>81</sup>. Less studied than *TFAP2A* |
| ***TOP1MT*** | 8 | Mitochondrial topoisomerase. Only indirectly affects cell fate | Over-expressed<br>Under-expressed | Relieves torsional strain by cutting and religating mitochondrial DNA during replication and transcription <sup>82</sup><br>Gatekeeper for cell fate decisions by controlling energy supply. |
| ***VAX2*** | 2 | Homeobox gene | Over-expressed | Translates positional information from *HOX* genes |
| ***GATA3*** | 10 | Master regulator of Th2 differentiation | Under-expressed | Upstream formation of gene enhancers. |
| ***MLPH*** | 2 | Part of the MYO5A-RAB27A trafficking pathway | Under-expressed | Decrease is a marker for high grade, poorly differentiated breast cancers. Tentatively implicated in cell differentiation pathways. |
| ***SPDEF*** | 6 | Downstream effector of NOTCH and IL-13/STAT6 signaling | Under-Expressed | Mucus production <sup>83</sup> Activated by morphogen-controlled pathways after progenitors have been committed to specific fates; participates in late-stage lineage specification. |
| ***XBP1*** | 22 | Gatekeeper for cells that require high secretory capacity. | Under-expressed | Activated B cells cannot become antibody secreting plasma cells without XBP1 |

Many genes with EBV-like methylation patterns are expressed exclusively in breast tumor epithelial cells and are incompatible with EBV infection restricted to infiltrating lymphocytes (TILs). Several are established cancer stem cell markers, including *ALDH1A3* (chr15), a retinoic acid enzyme marking breast epithelial cells and absent from TILs^32^, and *LGR4* (chr11), which is not expressed in immune or myeloid lineages. *SOX9* (chr17), a master regulator of basal lineage plasticity and stemness ^33,34^, is likewise not expressed in immune cells ^35^. *TCF7L1* (chr2), which enforces a pluripotent, non-differentiating state, is overwhelmingly epithelial and not expressed in lymphocytes ^36^. Core regulators *TFAP2A, OCT4*, and *SOX2* are not expressed in TILs; they normally cooperate to initiate exit from pluripotency ^37^; methylation of *TFAP2A* regulatory elements locks epithelial cells into self renewal.

Additional over expressed genes near EBV related methylation sites include *BMP2* (chr20) and *BMP4* (chr14), morphogens restricted to epithelial and stromal compartments, not TILs. *SMO* (chr7), a Hedgehog pathway receptor, is epithelial/stromal rather than lymphoid. Elevated *SPAG6* (chr10) does likely reflect immune infiltrate, whereas *CLDN3* (chr7) dysregulation compromises epithelial barriers ^38^ and is not expressed in lymphocytes.

Developmental homeobox genes *SIX2, SIX3,* and *VAX2* (chr2) are over expressed in breast tumors, but are absent in lymphocyte lineages, and single cell datasets consistently localize them to malignant epithelial clusters ^2..^ Within the *HOXC* cluster (chr12), *HOXC10* over expression originates from ER positive epithelial cells ^39^, while *HOTAIR* drives chromatin remodeling, *HER2* signaling, and estrogen responses, with no detectable expression in lymphocytes ^40^.

Several lineage specifying genes are under expressed, reflecting viral like effects on epithelial—not lymphoid—cells. *GATA3* (chr10), a luminal epithelial transcription factor marking basal-like tumors, not immune cells ^41^. Loss of *SPDEF* (chr6) impairs epithelial differentiation, shifting toward a basal, undifferentiated state, while *TFF3* (chr21) contributes to estrogen responsive luminal differentiation, not TILs.

Overall, the genes highlighted in Fig. 4C show that abnormal regulation of loci near EBV associated methylation sites primarily disrupts progenitor cell differentiation and promotes stem like cancer cell states. Their expression patterns are tumor intrinsic and incompatible with a model in which EBV-like methylation arises solely from infiltrating lymphocytes.

### Both ER-Negative and ER-positive breast cancers exhibited methylation at positions that parallel EBV-associated cancers

ER-positive tumors arise within the breast duct luminal lining, a compartment that contains well established stem-like cells. Basal cells occupy the outer duct layer and lack classical multipotent stem cells. Basal-like tumors, typically ER-negative, also originate in the duct lining, not from basal cells but from luminal progenitors that acquire a basal type phenotype^1^. ER-positivity is a strong surrogate for luminal breast cancers and ER-negativity reliably marks basal-like disease. Rare exceptions occur in both directions, but ER-subtype discordance in METABRIC is uncommon ^42^.

If EBV pushes luminal progenitors toward a basal program, then ER-negative and ER-positive tumors have a shared upstream mechanism, so they should both have methylation sites related to EBV-associated cancers. Fig. 5 shows this pattern: ER-negative tumors share 10–52% of their cis-methylation sites with EBV-associated cancers across 22 autosomes and ER-positive breast tumors show similar strong concordance with EBV-linked methylation patterns. Across both subtypes, 453 stem cell related genes were located near methylated regulatory elements shared with EBV-associated cancers; 296 in ER-positive tumors, 157 in ER-negatives and 113 in both. These results suggest that EBV drives or selects for stem cell states that can contribute to both basal-like (ER-) and luminal (ER+) breast tumors. Both stem cell states are luminal progenitor cells, but reprogrammed in different genomic backgrounds (such as *BRCA1* gene mutation status).

**Fig. 5.**
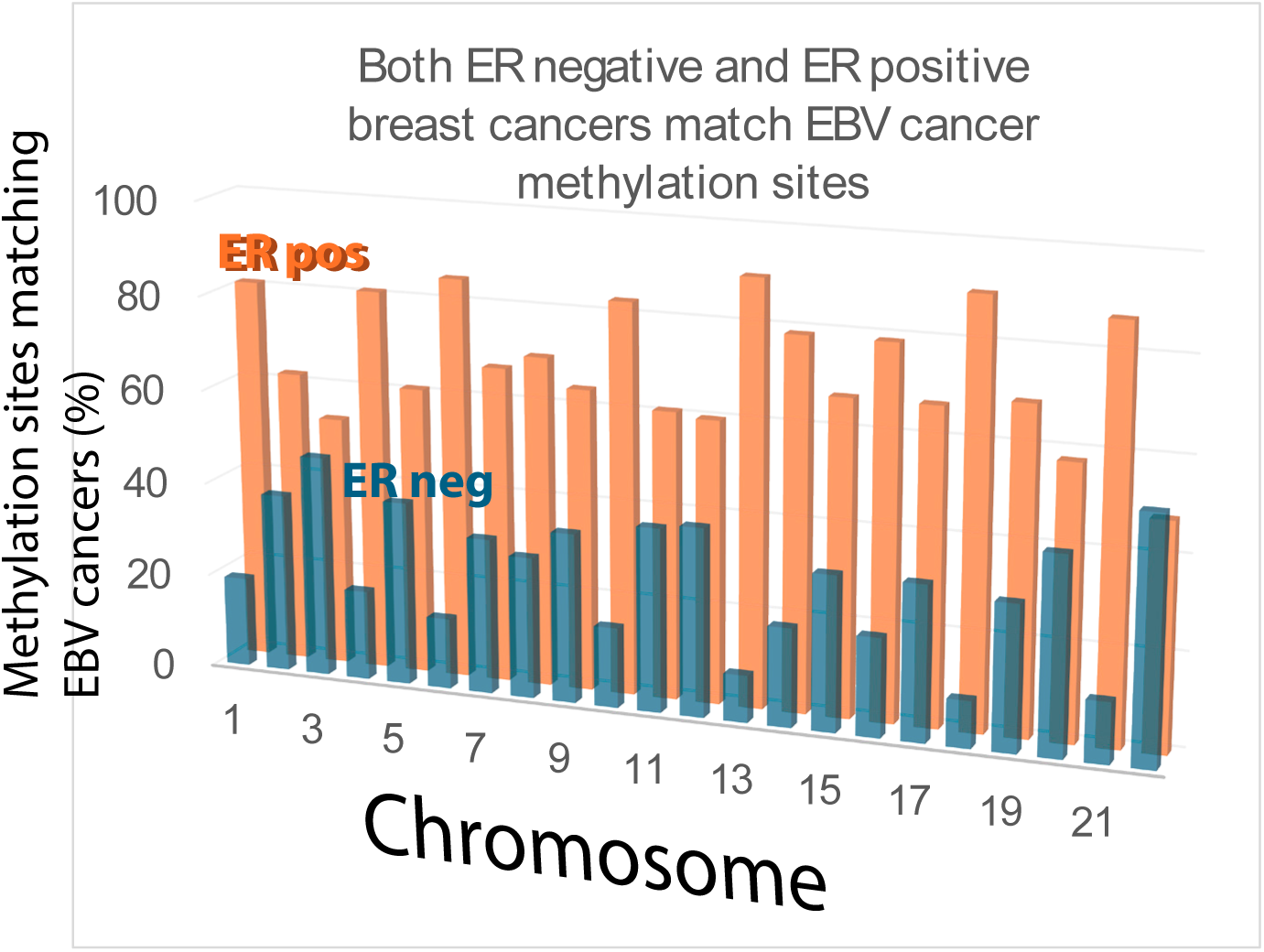
Both ER-negative (typically basal type) and ER-positive (typically luminal type) breast cancers have abnormal cis-methylation in sites that closely match cancers associated with EBV. Matches occur on all 22 autosomes and the shared sites are predominantly stem cell related genes.

### EBV related methylation scars in nonmalignant cells resemble breast cancer methylation patterns

To what extent do the results thus far apply to non-malignant cells with an active or a resolved past EBV infection? Specifically, are EBV induced methylation changes in normal cells related to those seen in EBV associated cancers?

In immortalized normal oral keratinocytes (NOKs), EBV infection induces proliferation, blocks differentiation^43^ and leaves persistent DNA methylation marks ^44^. These scars strongly correlated with breast cancer methylation (r=0.93, r^2^=0.87, p<0.0001), a result supported by Kolmogorov-Smirnov (p=0.89) and Spearman tests (p=0.88, CI=0.72-0.95). Chromosome level analysis reinforced this relationship. On chromosome 22, for example, methylation of *RAB36* control elements lies only 7 bp from a breast cancer methylation site—far closer than expected by chance (p = 0.012). Genome wide comparisons found 14 of 22 autosomes contained methylation scars near loci regulating stem cell related genes in breast cancer (Fig. 6). Aberrantly methylated gene control elements included:

**Fig. 6.**
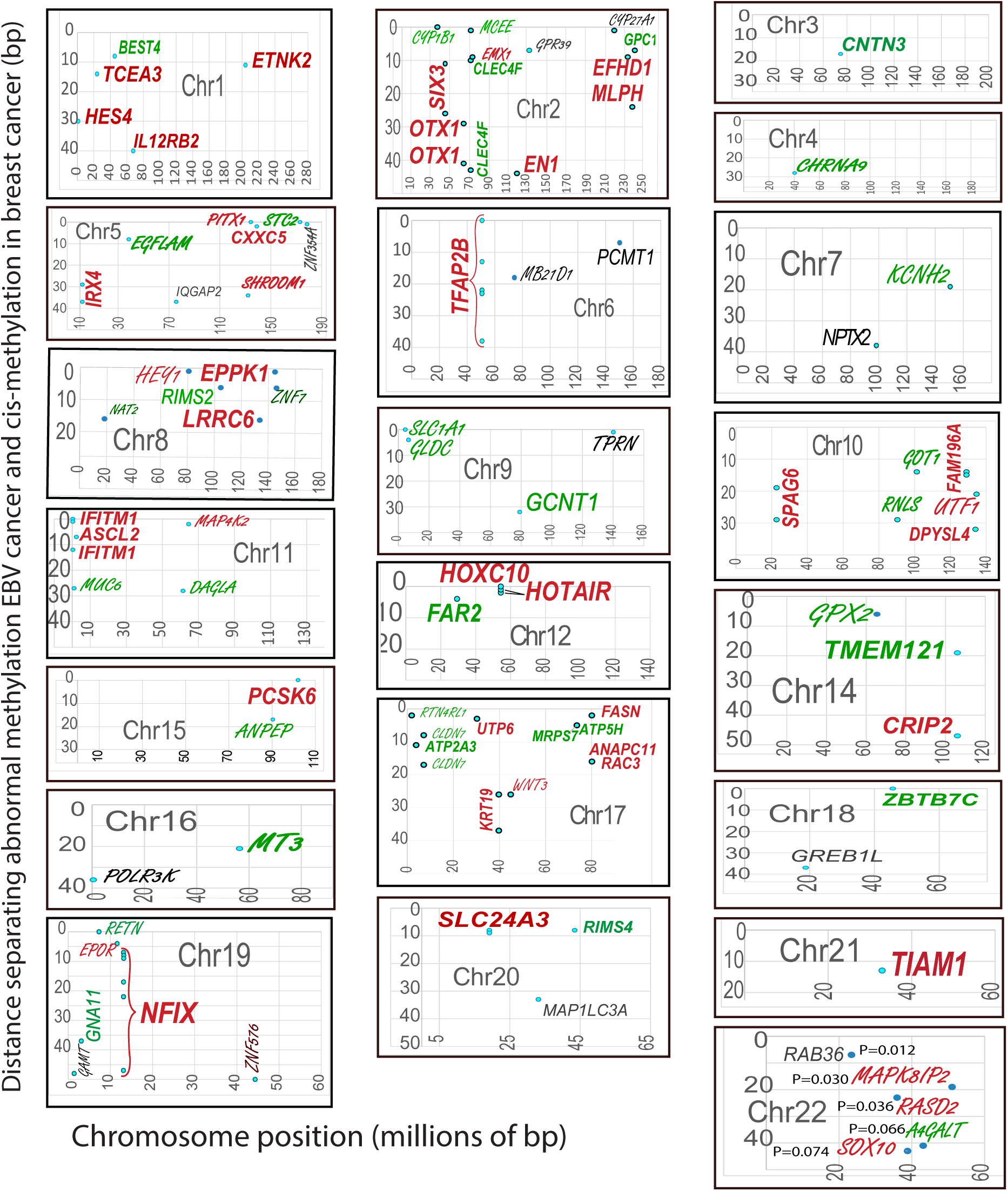
Gene-regulatory elements in breast cancer show abnormal cis-methylation at positions that also occur in non-malignant oral keratinocytes after recovery from EBV infection. NOK positions matching either NPC or BL were compared with abnormal cis-methylation in breast cancers. Most shared sites involve stem-cell–related gene controls (red) or microenvironment-related genes (green); other matched genes are shown in black. Genes methylated in EBV-exposed keratinocytes but not in breast cancers appear in script font. For chromosome 22, p-values show most matches are unlikely due to chance.

Chr2: 20/27 genes linked to stem cell regulation.

Chr12: *HOX*C and *HOTAIR*, mirroring breast cancer patterns.

Chr19: *NFIX* shared across breast cancer, EBV-associated cancer, and EBV infected NOKs.

Eight additional chromosomes contained newly identified methylation loci still connected to breast cancer stem cell pathways (Supplementary Table S2).

### Further control experiments

Skin basal cell carcinoma (BCC) has no known association with EBV and was used to test whether cancers unrelated to EBV share keratinocyte methylation changes. Nearly 7,000 hypermethylated BCC sites ^18^ were compared with abnormal methylation positions previously mapped in NOKs that had undergone EBV infection and were later cured. As shown in Fig. 7, overlap between BCC sites and EBV-associated keratinocyte sites ^40^ occurred less often than expected by chance. Within a 10 bp window, only one match was observed vs. 1.32 expected by chance (p = 0.73) and remained non-significant at 50 bp (p=0.27).

**Fig 7.**
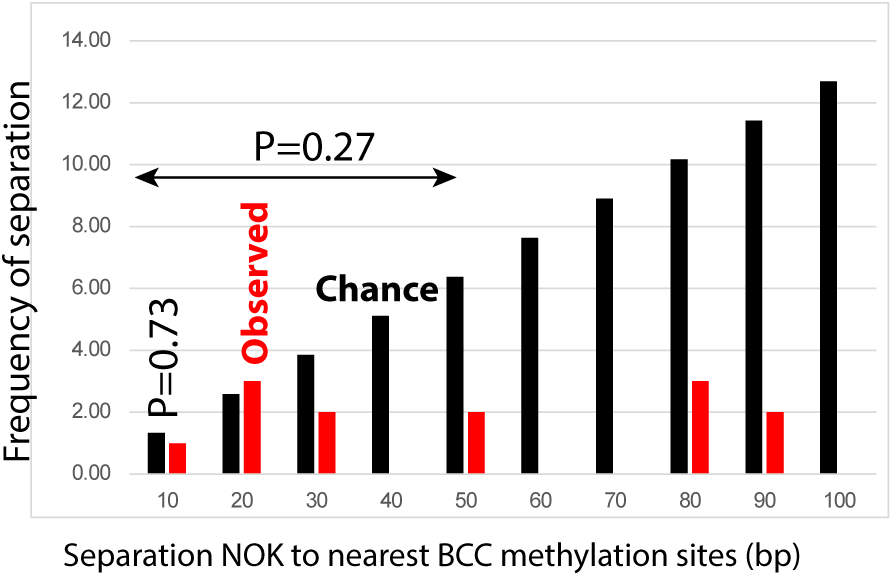
The frequency of methylated genes in normal oral keratinocytes that have recovered from an EBV infection (NOK)s within 50 bp of methylation sites in basal cell carcinoma of the skin is generally less than expected by chance.

These results indicate that methylation changes in EBV-infected NOKs differ from those in an EBV-unrelated cancer and are not random. Even after viral clearance, EBV infected keratinocytes retain distinct methylation marks, including some shared with malignant breast cells. These persistent “epigenetic scars,” suggest EBV induced methylation is difficult to fully erase and potentially dangerous.

When DNA replicates, newly copied strands are not methylated and depend on DNMT1 to copy the original pattern. Copying errors slowly accumulate over many cell divisions, producing an epigenetic replication clock. Because breast cancers have repli-cation-linked methylation ^12^, this clock could, hypothetically, explain similarities in breast and EBV cancers. To test this, sites where methylation increased in old cells vs. young cells ^45^ were compared with methylation sites in breast cancer. Only one CpG methylation site across four chromosomes agreed with breast cancer, indicating minimal contribution from repllication clock errors, consistent with published findings ^12^.

### Breast cancer dysregulates antiviral responses to EBV infection

Results thus far are compatible with two different origins for methylation concordance. One possibility is a resolved viral infection has left persistent regulatory scars more severe than in oral keratinocytes. Exogenous infection could induce a broad interferon response and widespread chromatin remodeling, followed by viral clearance and contraction to a restricted set of interferon stimulated genes (ISGs) that remain elevated due to sustained accessibility at antiviral loci. Persistent ISG expression, longterm repression of differentiation genes, partial erosion of retroelement silencing machinery, and altered transcriptional homeostasis would be seen. The past infection permanently remodeled chromatin and the epigenome despite no ongoing viral replication.

An alternative is the tumors themselves continuously drive a stem-like state. A cancer stem cell enriched signature arises from endogenous activation of innate immune pathways rather than exogenous viral infection. Derepression of endogenous transposons, cytosolic DNA or double stranded RNA generated by chromosomal instability, micronuclei rupture, and APOBEC mediated mutations could constitutively activate interferon signaling. This continuous signaling disrupts lineage specific enhancers and promotes stem-like cancer cell states.

Comparing *piRNA* distributions across chromosome 6 provided the first test of these alternative models. In the extended MHC region, *piRNA* clusters, *HLA* loci, endogenous retroviral fragments, and other transposon elements all occupy the same highly polymorphic block ^46^. Within this region, *piRNAs,* EBV-related and retrovirus-like sequences form an interspaced pattern resembling a CRISPR array, suggesting a structural principle of bacterial antiviral systems has been evolutionarily conserved in humans ^47^. *piRNA-PIWI* complexes recruit DNMT3 methyltransferases and other epigenetic machinery to silence these sequences ^48^ and shape chromatin around nearby MHC genes.

Because these antiviral and epigenetic functions depend on which PIWI proteins (PIWIL1-PIWIL4) load and execute piRNA activity, expression of the four proteins was examined in tumors. Across the four PIWIL paralogs, Fig. 8A shows a remarkable asymmetric pattern: *PIWIL2* and *PIWIL4* were predominantly and strongly over-expressed (z > 2), whereas *PIWIL1/3* were comparatively suppressed (z<-2) or largely dispensable. *PIWIL2/4* are the principal activated *piRNA*–epigenetic effectors in cancer; they suppress retroelements, dampen antiviral signaling, and stabilize stem cell–like chromatin states. *PIWIL4* expression showed a modest correlation with *SETDB1* (r=0.35), a histone methyl-transferase that coop-erates with *DNMT*s to silence endogenous retroviruses, modulate immune responses, and regulate pluripotency and cell fate. Given strong ISG induction, *PIWIL4–IFI16* correlation (r= 0.38) represents a bona fide antiviral/interferon response, compatible with exogenous viral infection, not endogenous retroelement driven viral mimicry.

**Fig. 8A.**
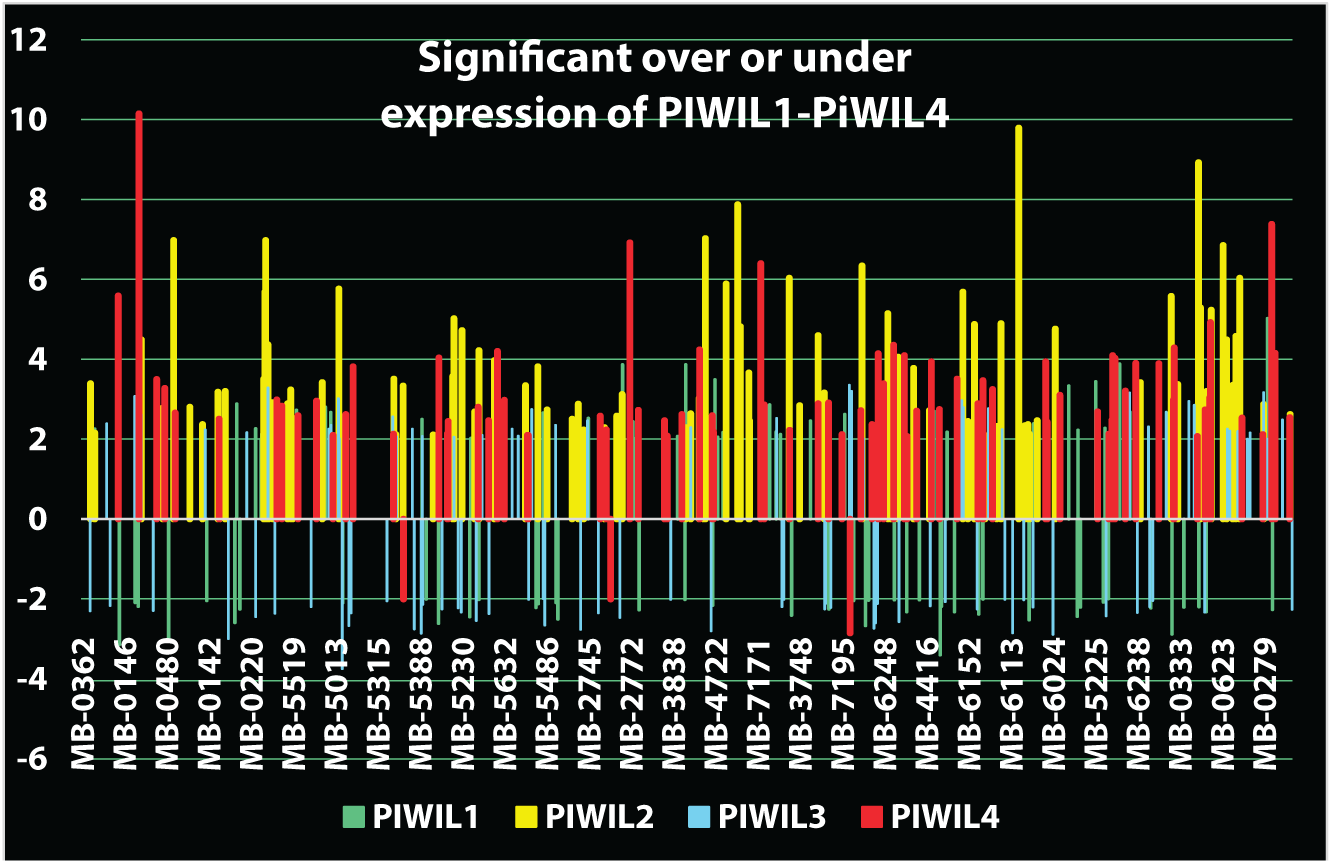
Dysregulation of the piRNA system in breast cancers. *PIWIL1-PIWIL4* genes with z>=|2| are shown. Overexpression occurs predominantly for *PIWIL2* and *PIWIL4*. *PIWIL1* and *PIWIL3*—linked to differentiation-associated *piRNA* pathways—are frequently sup-pressed, consistent with tumors avoiding developmental programs that oppose stemness or increase viral-mimic stress.

**Fig. 8B.**
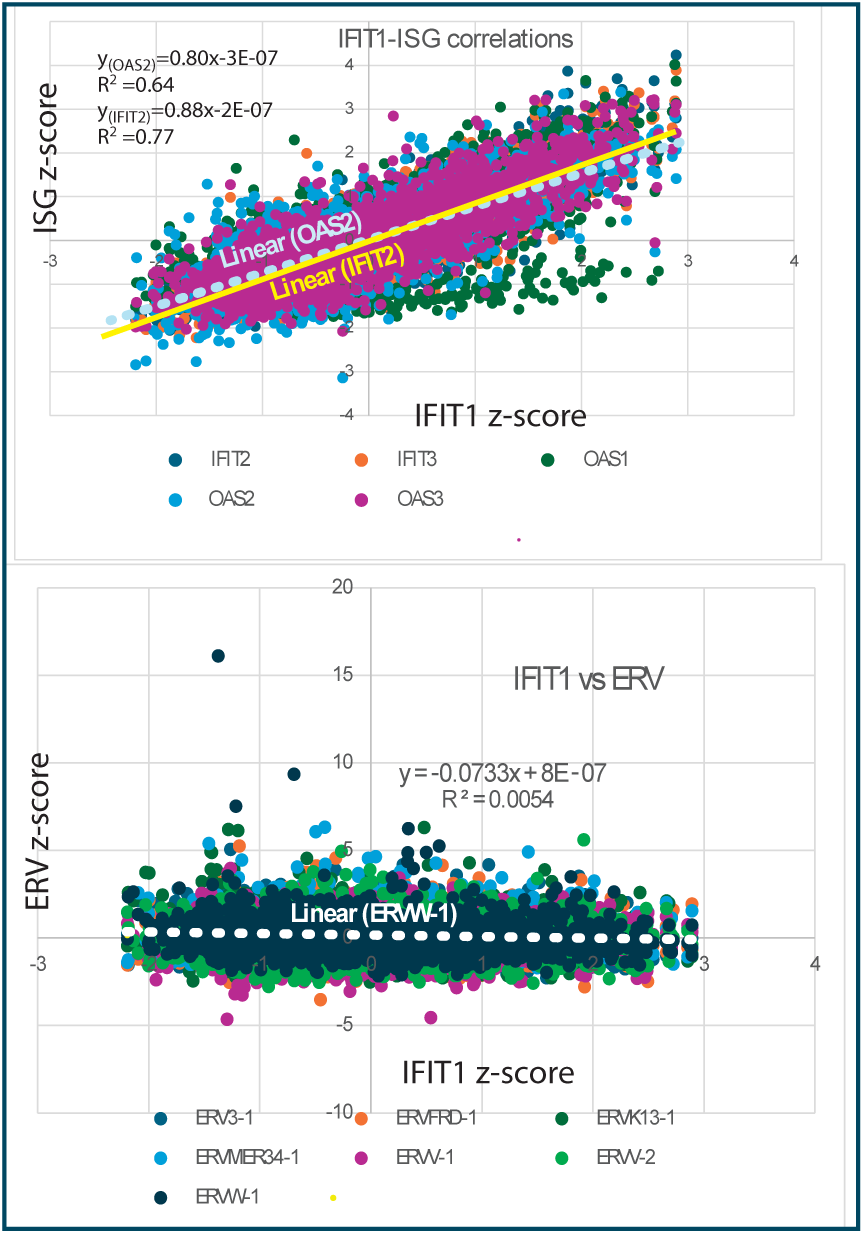
IFIT1 correlates strongly with other ISGs but has no coherent re-lationship to ERVs expressed in breast cancers

**Fig 8C.**
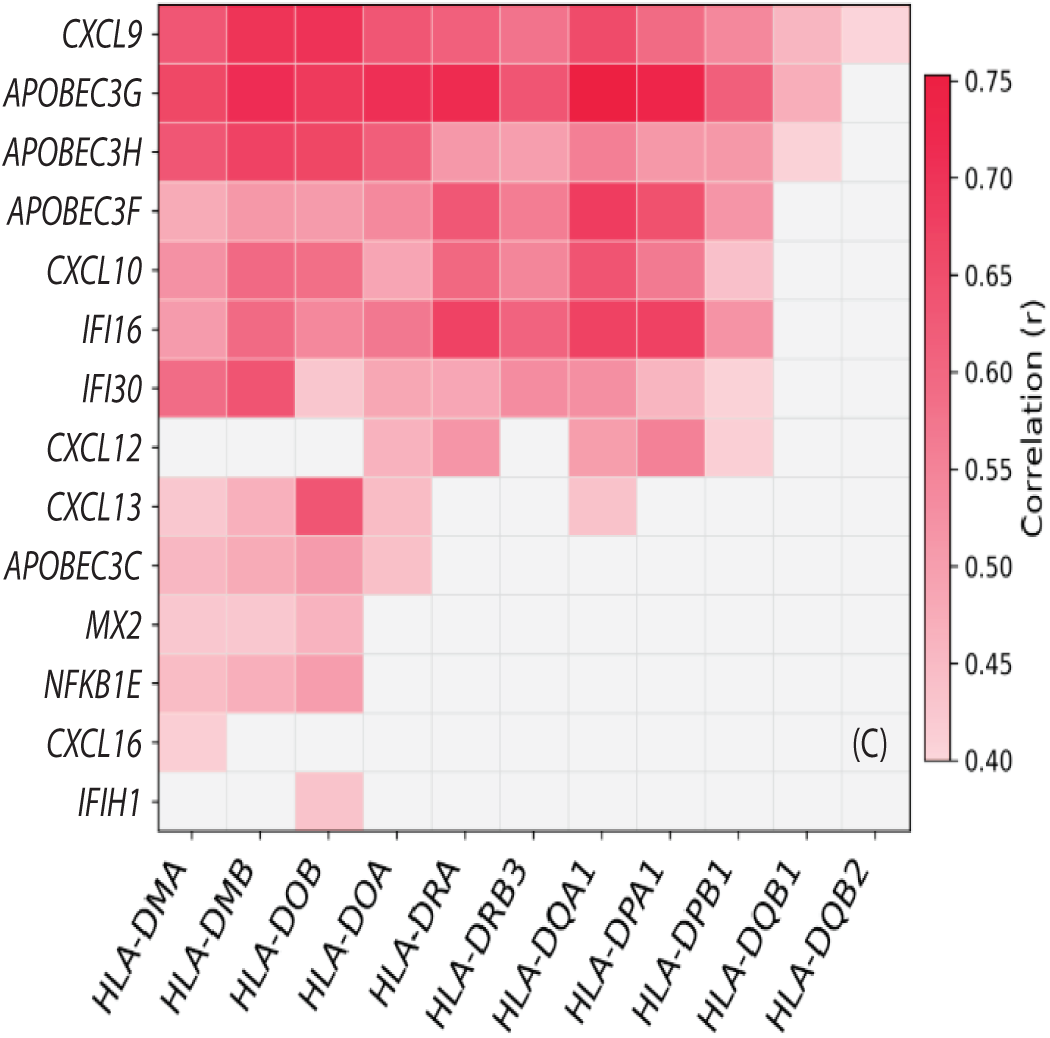
Interferon stimulated genes strongly correlate with HLA-D antigen presentation genes. These correla-tions indicate breast cancers have been responding to an exogenous infection and make molecular mimicry unlikely.

**Fig 8D.**
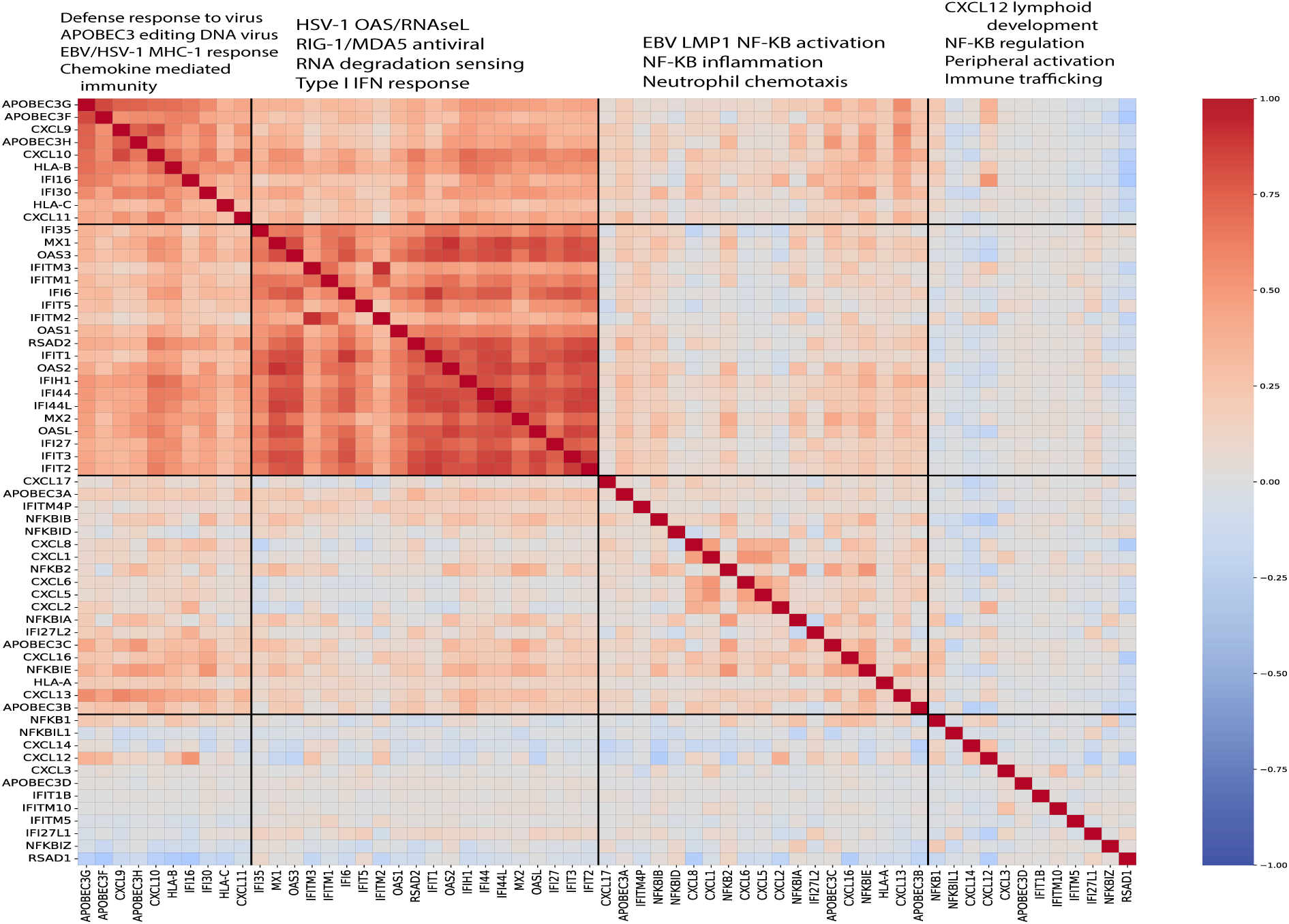
Four antiviral innate and adaptive immune pathways expressed in breast cancer typically engaged during viral infection. Abnormally expressed antiviral modules, especially modules 1 and 2, are co-expressed together. Alternate explanations not involving a virus have no support based on available medical information.

**Fig. 8E.**
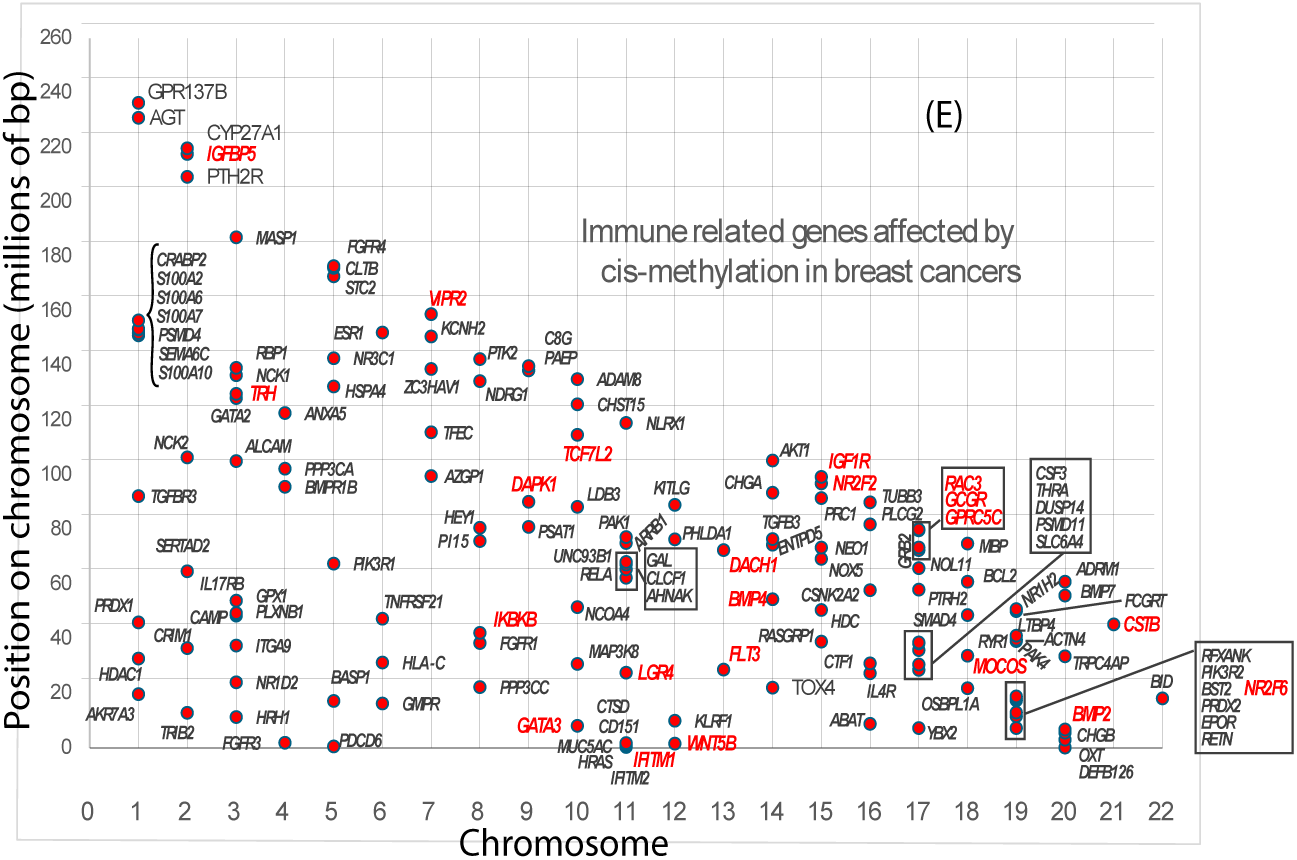
Immune response genes are affected by cis-methylation in breast cancer. The 4,012 breast cancer genes under cis-methylation con-trol include immune related genes on every autosome that often cluster together. Red text indicates stem cell related genes

**Fig 8F.**
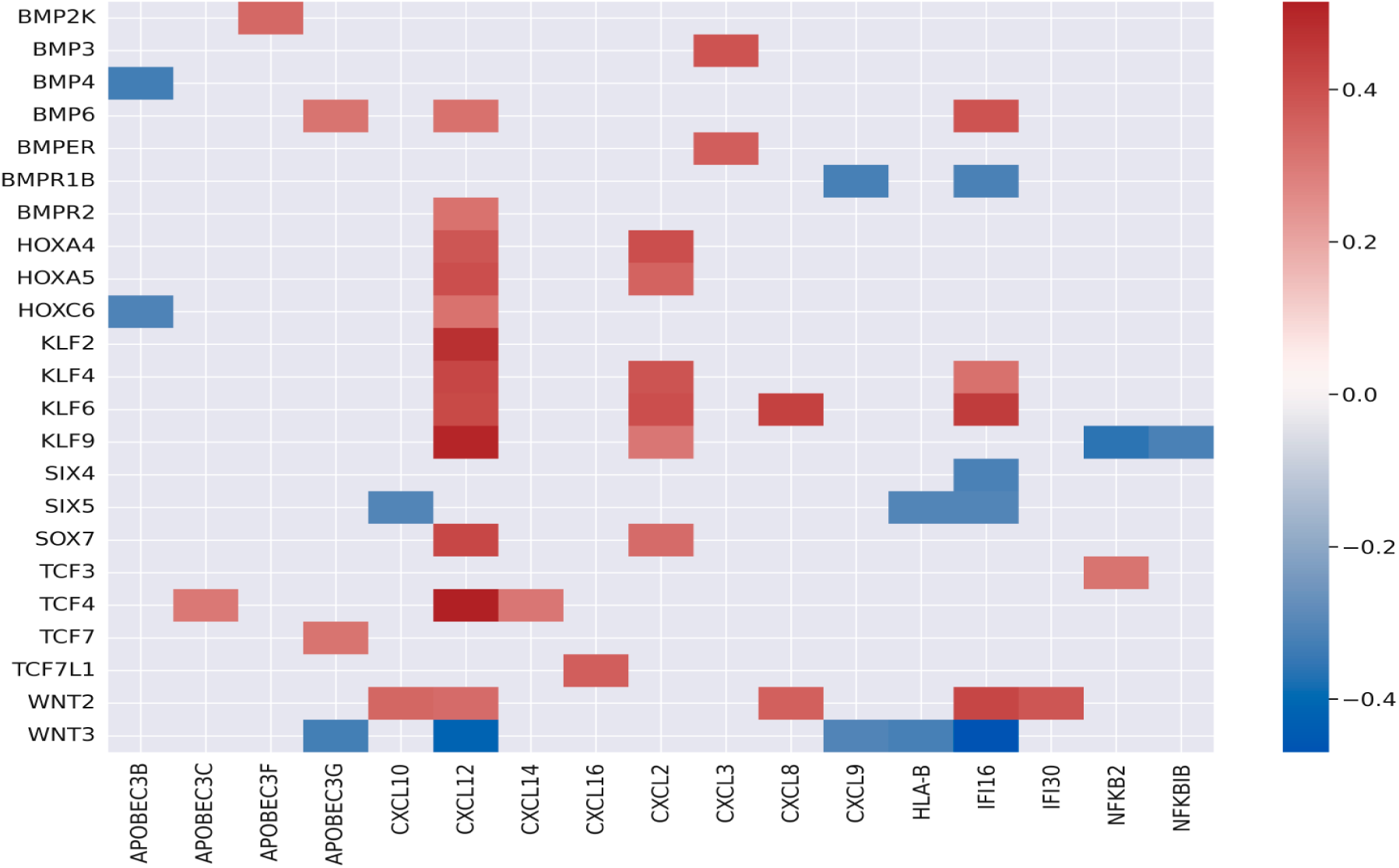
Four antiviral innate and adaptive immune pathways expressed in breast cancer typically engaged during viral infection. Abnormally expressed antiviral modules, especially modules 1 and 2, are co-expressed together. Alternate explanations not involving a virus have no support based on available medical information.

Fig. 8B illustrates the clear separation between exogenous antiviral interferon programs and endogenous retroelements (ERVs). When gene expression is compared as z-scores, exogenous ISG’s (*IFIT2, IFIT3, OAS1/2/3*) have a tightly coordinated linear relationship with *IFIT1*, with a slope near 0.9 with R²=0.77. These genes rise and fall together within a single, coherent interferon response. In contrast, ERV families show slopes and R² values near zero when plotted against *IFIT1.* Other ERV and HERV families are not detectably expressed. This pattern demonstrates that ERVs are transcriptionally silent or behave as background noise, without coupling to or stimulating the interferon pathway. Taken together, the data indicate that ISGs reflects a structured, exogenous virus-like response rather than endogenous viral mimicry driven by ERV derepression. ISG expression driven by other sources of chromosome instability are also unlikely.

Results in Fig. 8C support a response to infection on another level. Strong correlation was found between antigen presentation *HLA-D* genes and ISGs. In addition, TCGA data from >9,000 tumors revealed 65 differentially methylated *HLA* genes across the *D, G, J,* and *L* loci, including 11 enhancers. Thus, antigen presentation pathways normally stabilized by piRNA-guided mechanisms become dysregulated in breast cancer. Dysregulation involves methylation dependent processes linked to interferon^49^ or extensive fragmentation of the *HLA r*egion ^30,47^.

The next analysis examined whether these and other biomarkers were coordinated as would be expected for a response to an infection. The heatmap in Fig. 8D pictures representatives of pathways typically induced by interferon or by viral infection, including well described elements of herpesvirus biology.

**Module 1** – Cytotoxic/IFNγ program, T-cell infiltration: *APOBEC3* family members (*A3F/G/H*), *CXCL*9/10/11, *HLA* class I genes, and the DNA virus sensor *IFI16*. These genes respectively mark antiviral DNA/RNA editing ^50^, Th1 chemokine mediated effector recruitment, CD8⁺ T-cell antigen presentation, and nuclear sensing of viral genomes/ episomes ^51^.

**Module 2** – Type I interferon (ISG) genes suggesting antiviral response: *OAS1/2/3, MX1/2, IFITs,* and *IFIH1/MDA5, IFIs.* Strong internal correlations suggest active IFNα/β signaling and dsRNA sensor engagement. Although EBV is a DNA virus, its EBER RNAs can activate *TLR3/PKR/MDA5* pathways and induce ISGs ^52^

**Module 3** – NF-κB–linked inflammatory chemokines: *NF-kB1/2, IκB* regulators, and *CXCL*1/2/5/8/13, consistent with *NF-κB* activation. EBV LMP1 is a potent, constitutive activator of *NF-κB* pathways, providing a mechanistic basis ^53,54^

**Module 4** – Stromal/tissue remodeling microenvironment: *CXCL12, CXCL14, NF-kBIL1, RSAD1*, and *IFI27L1*, reflecting chemokine guided organization and stromal activation often observed in chronically inflamed, virally affected tissues. *CXCL12/CXCR4* and *CXCL14* have established roles in tissue organization and chemokines that can accompany persistent viral states ^55^.

Fig. 8D shows strong correlations within and among certain modules. The correlations within module 1 suggest TILs are being recruited. Module1-module 2 have strong correlations not attributed to an exogenous virus. The general coordination trend varies among patients and is not guaranteed for any individual. High expression of *CXCL9/10, HLA-D/E/F,* and *APOBEC3* indicate a T-cell inflamed tumor microenvironment, characteristic of basal-like tumors. In contrast, luminal, ER-driven tumors have low expression of *APOBEC3, HLA* genes, and low TILs and high estrogen signaling ^56^. NK inhibitory ligands HLA-E ^57^, and HLA-F ^58^, which are upregulated during EBV infection, are also upregulated in breast cancers. These correlations reinforce the case for an ongoing or lingering response to virus.

### The antiviral response in breast cancers affects differentiation of progeni-tor cells

*PIWIL4* expression correlates negatively with *FOXA1* (r=-0.5) and *GATA3* (r=-0.43) indicating that activation of germline-like epigenetic programs emerges as luminal identity wanes, consistent with a shift toward a more plastic, less differentiated state.

HLA and interferon stimulation effects on stem cell differentiation were then assessed. Of 4,012 autosomal cis-methylated genes ^12^, 154 also appeared in a separate list ^59^ of 1,480 immunity genes (Fig. 8E). About 22 stem cell–associated genes were within this intersection. Incorporating additional components of shared stem cell differentiation pathways ^60^ increased the total to at least 50. These results suggest that EBV-like methylation patterns extend into stem cell regulatory programs integral to effective immune responses.

The next step was to assess whether the overall coordinated expression of gene modules influences cell lineage programs in breast cancer. As shown in Fig 8F, genes representing major developmental pathways (*HOX, WNT, BMP, SOX, TCF, KLF, SIX, VAX*) correlate strongly with interferon stimulated and immune response genes (*CXCL, IFI, APOBEC3, HLA, NF-kB*). The genes *CXCL2, 3, 6, 8* clustered with developmental and stromal organization genes, whereas *CXCL*9 was strongly anti-correlated with *WNT3* and *BMPR1B*, consistent with an *IFNγ/STAT1*driven inflammatory module. *CXCL12* showed a mixed but predominantly stromal pattern—negative vs *WNT3* yet positive vs *WNT2, TCF4, SOX7, KLF2/4/6/9, BMPR2, and BMP5*—aligning with *CXCL12* niche forming and vascular patterning ^65^. *APOBEC3B* was negatively correlated with *HOXC6* and *BMP4*, and *HLAB* was negatively correlated with *WNT3* and *SIX5*, further supporting the antagonism between interferon linked genes and developmental programs. *NF-kB2* and *NF-kBIB* were negatively associated with *KLF9,* although *NF-kB2* remained positively correlated with *TCF3*, indicating selective coupling of *NFκB* signaling to specific developmental regulators. In general, these patterns have inter-feron and *NFκB*–associated genes opposing core developmental pathways, while *CXCL12* and *CXCL2/3/6/8* define a stromal/developmental module. These results suggest that infection delays lineage commitment.

An alternative explanation is that an endogenous *DUX4-*like program derepresses ERVs, producing the *APOBEC* and interferon related signatures observed. However, across 1,980 breast cancers, *DUX4* itself was not expressed, and the nine detectable *DUX4* paralogs or pseudogenes were almost uniformly under expressed. Because DUX family proteins function exclusively as transcriptional activators ^61^, their reduced expression cannot generate a *DUX*-like ERV or interferon program. Accordingly, these paralogs showed reduced expression in 85% of tumor–gene measurements (15,225 of 17,820), arguing strongly against an active *DUX*-like transcriptional program as the source of the observed signatures.

## Discussion

Breast cancers display methylation changes in the same stem cell regulatory elements altered in EBV-cancers and even in cells that have cleared EBV. These cell states are selected for because they are permissive to viral replication and they contribute to tumor heterogeneity and plasticity. Selection occurs in both ER-positive and ER-negative breast cancers, which then produce luminal or basal-type breast cancers. Comparisons to non-EBV skin carcinoma, randomized sites, retroelements, TILs, *DUX4*, and the replication clock confirm the specificity of EBV-linked methylation associations.

EBV broadly alters accessibility and expression of lineage defining transcription factors, including *HOX* genes and their interacting morphogen pathways. Aberrant *HOX* methylation can disrupt tissue patterning, enhance oncogenic activity, impair tumor suppression, and increase sensitivity to environmental carcinogens.

Methylation patterns and compromised genome safeguards are characteristic evidence of prior EBV infection, even after viral transcripts disappear ^47^. The Fig. 8 signatures—*APOBEC3* activity, *IFI16* sensing, and LMP1like *NFκB*—most closely match a DNA virus response (e.g., EBV, HSV1). However RNA viruses have some overlap as expected.

Correlations between progenitor cell regulators (*HOX, WNT, BMP, SOX, TCF, KLF, SIX, VAX*) and interferon stimulated immune genes (*IFI, IFIT, APOBEC3, CXCL, HLA, NF-κB*) further support viral involvement. Viral nucleic acids detected by *RIGI, MDA5, TLRs*, or *cGAS–STING* trigger type I/III interferon programs that reshape cell state pathways, affecting differentiation, growth arrest, and stress repair pathways.

Potential non-viral explanations for the correlations observed are bacterial sepsis, autoimmunity, allergic reactions, transplant rejection, chronic inflammatory disease, and medication effects. Genomic studies of breast cancer typically do not evaluate these non-viral comorbidities, but large observational studies show that they are rare and contribute minimally to breast cancer prevalence ^62^. Although the pattern is not uniquely diagnostic, viral infection remains the simplest explanation. Many stimuli that activate interferon and *NFκB* pathways (inflammation, DNA damage, replication stress, cytokines, retroelement derepression, or microenvironmental cues) are downstream from viral activity. Consequently, transcriptional, and epigenetic signatures often labeled as non-viral may instead represent remnants from a prior or occult viral event, blurring the distinction between active infection and its long lived molecular footprint.

The tight coupling among *CXCL, APOBEC3*, and broader ISGs in breast cancers suggests a shared upstream inflammatory driver. This pattern resembles the interferon dominant program characteristic of EBV-associated malignancies, where EBV robustly induces *CXCL*10, activates *RIGI* and c*GAS–STING* signaling, and upregulates *APOBEC3* family members. The pattern is not reproduced in all cancers, suggesting alternatives or that it often fails. Although EBV status was not assessed, EBV infects essentially all adults and leaves durable immune and epigenetic alterations long after becoming undetectable.

Mechanistically, trans-regulation readily explains coordinated *CXCL*10 (chr4) and *APOBEC3* (chr22) expression: interferon activated *JAK–STAT* signaling induces *STAT1/IRF1,* which co-regulates both gene sets. Pronounced *APOBEC3* locus hypomethylation and over-expression in some tumors is therefore most consistent with secondary epigenetic consequences of sustained interferon signaling rather than primary cis-regulatory changes. Together, these data indicate activation of a high interferon state that phenocopies EBV-associated inflammation. This program explains *CXCL–APOBEC3* co-induction, even in the absence of direct viral detection. Similar methylation patterns in other EBV-linked cancers support a shared epigenetic mechanism. EBV-driven reprogramming is well documented in NPC. NPC shows widespread hypermethylation, altered DNMT3A/3B activity and resistance to demethylation, altering chromatin accessibility. BL also exhibits EBV-associated epigenetic reprogramming. Both NPC and BL can occur in EBV negative forms, consistent with EBV initiated epigenetic changes persisting even after virus becomes undetectable.

EBV repeatedly targets the *CEBPA/CEBPB* differentiation axis, through epigenetic marks on infected stem cells—pushing EBV-cancers ^63,64^ toward stem-like or dedifferentiated states ^65^. *CEPB* regulatory regions show EBV-like methylation in breast cancers.

Although there are many potential confounders, epidemiologic studies suggest that EBV is significantly more common in breast cancers than in control tissues ^66^. The present findings align more strongly with EBV exposure increasing breast cancer risk than with breast cancers merely becoming more permissive to infection. Even after viral clearance, persistent methylation and genome damage ^47^ maintain a pro-tumorigenic state, decoupling risk from ongoing infection. Breast cancers exhibit comparable epigenetic and genomic alterations at many of the same sites as malignancies with established EBV connections. Together, these data support a model in which transient EBV infection leaves lasting molecular damage that elevates breast cancer susceptibility. EBV exposure perturbs stem cell hierarchies ^67^, bypasses genome stability safeguards ^47^, and interacts with carcinogens to influence whether infection resolves or progresses toward breast cancer.

These findings also underscore the susceptibility of stem cell populations to virus induced epigenetic reprogramming, reinforcing the broader value of vaccination against oncogenic viruses and highlighting the need for rigorous epigenetic quality control in stem cell–based therapies.

## Resource Availability

Datasets used are well-documented and freely available from original sources or the author on reasonable request.

## Methods

### Breast cancer patient data

Breast cancer data came from 1904 females with primary breast cancers, mainly from the METABRIC cohort. Original publications ^30^, cBioPortal, and Kaggle are freely available sources genome data. Briefly, 1459 tumors were ER-positive and 445 were ER negative. Grades 1, 2, and 3 were in 165, 740, and 927 patients, respectively. Most cancers (1454) were ductal. Patient ages were ∼22 to 96, with most (1200/1904) ages 47-72. Original publications included clinical descriptions, tissue collection procedures, gene expression profiles, copy number aberrations, and point mutations ^30,68^. Stage, tumor size, lymph node status, PAM50 genes, and driver genes were also available.

### Breast cancer methylation data

DNA methylation data were from published whole genome reduced representation bisulfite sequencing of 1,538 breast tumors and 244 adjacent normal tissues ^12^. The study screened promoters for Pearson correlations with methylation sites located on the same (“cis”) chromosome. Broad (“trans”) methylation effects influencing genes on multiple chromosomes were excluded. Assigned cis-acting methylation sites showed the strongest, most negative correlation with their corresponding genes across more than 50 tumors (FDR < 0.05). Additional methylation and expression data were downloaded from TCGA using the BIOLINKS package in R through the Posit/RStudio interface.

### NPC patients and methylation data

Comparisons between methylation patterns in breast cancer and NPC used independently published datasets ^9^. NPC methylation data were derived from whole genome bisulfite sequencing of biopsies from 15 sporadic NPC patients and 9 matched non tumor adjacent tissues. All tumors were primary NPCs (stage II, n=3; stage III, n=7; stage IV, n=5). Patients ranged from 38–82 years of age (13 males, 2 females), and 11 were survivors.

Differential methylation in NPC was the average difference between NPC and matched normal tissue. A total of 16,910 differentially methylated regions (DMRs) showed absolute methylation differences ≥0.2, and 6036 were <500 bp long with average differential methylation of-0.336 to 0.463. Three NPC samples were globally hypomethylated and 12 globally hypermethylated. Whole genome bisulfite sequence results had been validated in an independent cohort of 48 NPC patients ^9^.

### BL patients and methylation data

Comparative data for Burkitt lymphoma (BL) were obtained from a published cohort of 13 patients (ages 3–18; 11 males, 2 female) ^15^. All tumors harbored *MYC* breaks with IG–MYC translocations and exhibited differential hypermethylation correlated with gene expression. More than two-thirds of the DMRs were shorter than 500 bp, with lengths varying from 8 to 5,160 bp. Reference DNA was from non-neoplastic germinal center B-cells from tonsils of four donors (ages 13–30). The present analysis used 42,381 hypermethylated DMRs across the 22 autosomes. Average methylation levels in germinal center B-cell DNA were subtracted from corresponding BL values, and only regions showing ≥20% differential methylation were retained,

### Transiently infected non-malignant cell data

Published source data^40^ came from hTERT immortalized NOKs infected by 24 h coculture with anti-IgG–induced Akata BL cells. Following B-cell removal, infected cells underwent 10 passages of antibiotic selection, followed by 10 passages without selection to obtain transiently infected populations. Single cell clones were isolated by flow cytometric sorting; EBER in situ hybridization confirmed EBV status. Uninfected parental NOKs and plasmid vector controls were cultured in parallel, and all clones were authenticated by DNA fingerprinting. Methylation status came from reduced representation bisulfite sequencing^40^.

### Skin basal cell carcinoma methylation data

Basal cell carcinoma data (BCC) came from 16 BCC biopsy specimens (11 women and 5 men, ages 55-89). The samples were 8 nodular and 8 sclerodermiform BCCs ^14^. Tumor sites included face, shoulder, head, ear, lip, nose, and pectoral areas. Whole genome methylation profiling was with Infinium MethylationEPIC BeadChips.

### Calculations of matching abnormally methylated positions

Distances between abnormally methylated positions in breast cancer and EBV associated cancers (NPC or BL) were computed as the minimum absolute difference between the start, end, and midpoint coordinates of each breast cancer cis hypermethylated segment and coordinates of EBV associated DMRs. A strict cutoff of ≥20% difference from normal cell methylation was applied across the full length of each segment. An example of a typical spreadsheet formula used for minimum distance calculation is as follows. =MIN(ABS(D2-$K$2:$M$1532),ABS(E2-$K$2:$M$1532),ABS(F2-$K$2:$M$1532)), where columns D through F contain breast cancer cis-hypermethylation positions and columns K through M contain differential hypermethylation positions in EBV cancer (either NPC or BL). The Poisson distribution model estimated the probability of a given number of rare, independent events occurring in a fixed space or time; p-values were then: P(X≥N)=1-P(X<N)=1-∑_(k=0)^(k=N-1) (e^(-λ) λ^k)/k! (where λ= number of matches expected from random events and N=observed count in the data). Poisson distributions were evaluated using a Python 3.13 script.

For control comparisons between breast cancer and skin basal cell carcinoma, only sites with β ≥ 0.5 were included, where β is the fraction of methylated reads relative to total methylated plus unmethylated reads. To evaluate potential artifacts from unusually long segments, control differential methylation intervals were truncated at 1000 bp; this adjustment produced only minor changes in the results. All genomic coordinates were taken directly from the original publications and map to GRCh37/hg19 human genome assembly. Genome positions were used directly from the original publications and not corrected or manipulated in any way.

### Calculation of frequency of close agreement

Frequencies of agreement between breast cancer and EBV cancer were calculated in Excel after normalizing the numbers of NPC differential methylation sites to the same total numbers of cancers as BL data (15 NPC vs. 13 BL).

### Relationships to stem cells and stem cell microenvironments

Stem cell associated genes were defined as those essential for self-renewal, asymmetric division, terminal differentiation from a pluripotent state, or regulation of these processes. Traditionally, “stem cell genes” refers to regulatory factors—such as *OCT4*, NANOG, or *SOX2*—that maintain pluripotency and self-renewal. In the present work, morphogens and other transcription factors that govern cell-identity specification and lineage programs were also included.

Approximately 350 genes with abnormal methylation within <50 bp of matched breast and EBV cancer or EBV infected non-malignant cells were manually curated for function. Microsoft Copilot and Copilot Pro were used and produced only infrequent errors so all outputs were manually verified against literature sources. Each breast cancer gene methylated within <50 bp of an EBV associated methylation site was evaluated for links to stem cell biology: regulation of stem cell gene expression, lineage determination, or use as a stem cell marker. A curated stem cell database ^76^ was regularly consulted. A gene was classified as stem cell–related only when supported by a credible reference.

Parallel searches examined whether shared breast cancer–EBV loci influenced the stem cell microenvironment. Genes affecting the microenvironment include those encoding extracellular matrix components, transport and structural factors, and other modulators that shape external or internal conditions affecting cellular responses. Genes were categorized as “Other” when their functions affected stem cell processes only indirectly.

### Breast cancer gene expression data

Breast cancer gene expression data came from the METABRIC study ^25^, which used the Illumina HT12 v3 microarray. Genes with expression z-scores ≥2 standard deviations from the mean (p < 0.05) were identified and compared with stem cell genes located within 50 bp of methylation sites associated with EBV related cancers.

### Calculation of overlap in immune related methylation positions in breast cancers vs methylation position in EBV cancers

Methylated CpG islands or map positions fall into methylation clusters that select for the immune cell infiltrate in breast cancers ^77^. Illumina IDs were converted into map position and island location. Minimum distances between these positions and abnormally methylated positions in NPC and BL cancers were then calculated. An Excel formula determined whether an EBV cancer methylation position in cell K2 was within the start to end range for any of three sets of methylation positions in CGI regions for breast cancer immune prognosis genes (in rows 2-4 across columns AB to AF). =IF(OR(AND(K2 >= MIN($AB$2,$AC$2,$AD$2,$AE$2,$AF$2) - 50, K2 <= MAX($AB$2,$AC$2, $AD$2,$AE$2,$AF$2) + 50), AND(K2 >= MIN($AB$3,$AC$3,$AD$3,$AE$3,$AF$3) - 50, K2 <= MAX($AB$3,$AC$3,$AD$3,$AE$3,$AF$3) + 50), AND(K2 >= MIN($AB$4,$AC$4,$AD$4,$AE$4,$AF$4) - 50, K2 <= MAX($AB$4,$AC$4,$AD$4,$AE$4,$AF$4) + 50)),K2, “”)

### Comparisons of distributions

Normality was assessed using StatsDirect. Hypermethylation distributions consistently deviated from normal and were evaluated using Mann–Whitney nonparametric tests. Correlations between breast cancer and EBV-associated epigenetic modifications were identified by highly powered linear regression. Kolmogorov–Smirnov and Spearman’s rank tests provided additional nonparametric comparisons. Frequency plots were generated to visualize and verify the distribution of coincident regions.

### Window sizes to compare methylation sites in breast and EBV cancers

Differential methylation sites in breast cancer and EBV-associated cancers were compared using windows <50 bp, consistent with the minimum length of enhancer elements (50–1500 bp). Three 50bp windows (start, middle, end) were applied to regulatory regions to restrict comparisons to <200 bp and capture only the shortest shared sequences within larger hypermethylated domains.

To ensure clearer interpretation, breast cancer methylation sites had an upper limit <50 bp of methylation positions in EBV cancers. To assess the stringency of the <50bp window, distances were quantified between methylation sites and transcription start sites (TSS). Methylated-site frequencies were available from –500 kb to +500 kb ^7^ around eight paired gene TSSs using pooled chromosome 1–22 data. Distances between breast cancer–specific hypermethylated sites and TSSs were then evaluated. Of 4,012 methylation sites, 927 were within 1 kb of a TSS, 261 within 500 bp, one within 200 bp, and none within 50 bp. Because no sites occurred within the <50bp range, the 50bp upper limit was considered stringent and used for all comparisons.

## Key Resources Table

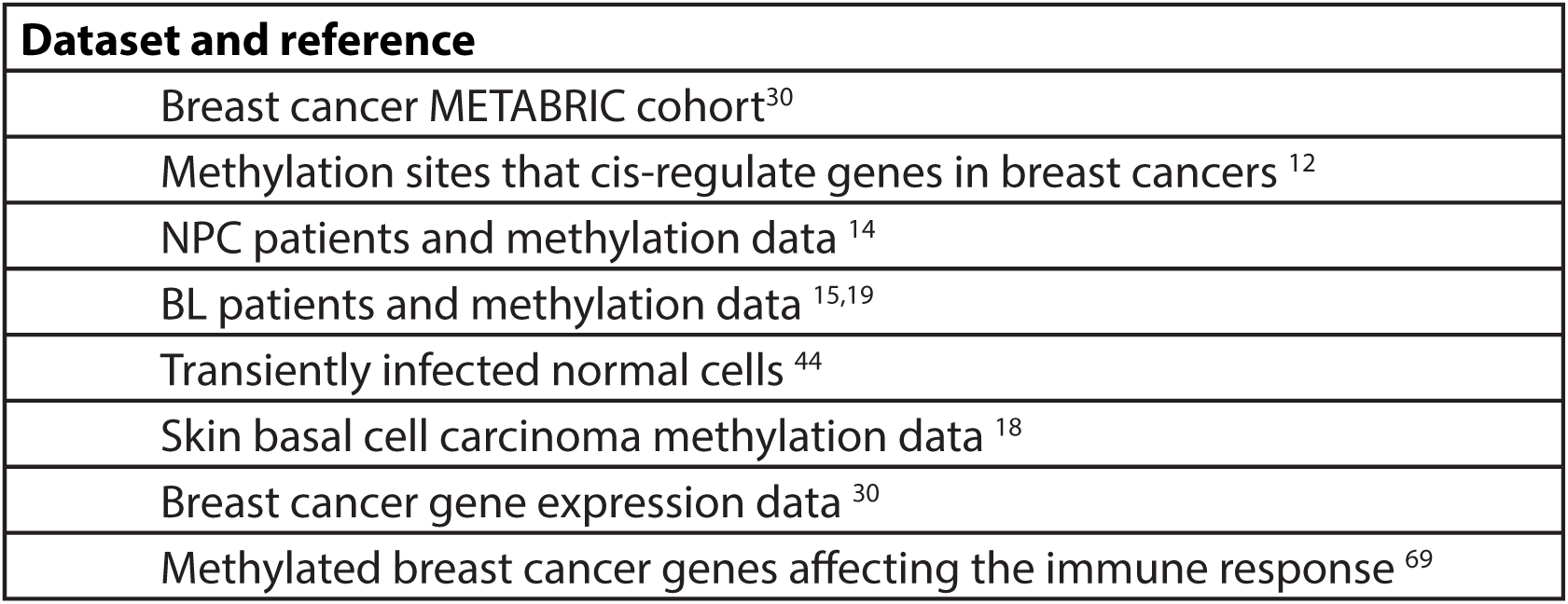

| Dataset and reference |
| --- |
| Breast cancer METABRIC cohort<sup>30</sup> |
| Methylation sites that cis-regulate genes in breast cancers<sup>12</sup> |
| NPC patients and methylation data<sup>14</sup> |
| BL patients and methylation data<sup>15,19</sup> |
| Transiently infected normal cells<sup>44</sup> |
| Skin basal cell carcinoma methylation data<sup>18</sup> |
| Breast cancer gene expression data<sup>30</sup> |
| Methylated breast cancer genes affecting the immune response<sup>69</sup> |

