## Supplemental Table S2 for "Epstein-Barr virus induced epigenetic reprogramming drives cancer stem cell emergence in breast cancer"

***Supplementary Table S2 Functions of gene control sites methylated in non-malignant oral keratinocytes that are not methylated in the breast cancer cohort***

| **Chromosome** | **Gene** | **Class** | **Encoded Function** |
|---|---|---|---|
| 2 | *CYP1B1* | Microenvironment | Detoxification |
| 2 | *CYP27A1* | Other | Cholesterol metabolism |
| 2 | *EMX1* | Stem cell | Stem cell self renewal and proliferation |
| 2 | *GPR39* | Microenvironment | Zinc homeostasis |
| 2 | *MCEE* | Microenvironment | Metabolic enzyme essential for energy production in mitochondria |
| 4 | *CHRNA9* | Microenvironment | Ion channel receptor for acetyl choline expressed in stem cells |
| 5 | *EGFLAM* | Microenvironment | Extracellular matrix organization |
| 5 | *STC2* | Microenvironment | Calcium phosphate homeostasis |
| 5 | *ZNF354A* | Other | Transcription regulator cancer biomarker |
| 6 | *MB21D1* | Other | Antiviral defense, DNA damage sensor |
| 7 | *KCNH2* | Microenvironment | Voltage gated potassium channel |
| 8 | *HEY1* | Stem cell | Downstream target of Notch signaling pathway, regulates self renewal and differentiation |
| 8 | *NAT2* | Other | Drug metabolism and detoxification |
| 8 | *ZNF1* | Other | Transcription regulation |
| 9 | *GLDC* | Microenvironment | Glycine catabolism, redox balance |
| 9 | *SLC1A1* | Microenvironment | Glutamate transporter essential for neural stem cell self-renewal [1] |
| 9 | *TPRN* | Other | Inner ear function, cell structure and function |
| 10 | *GOT1* | Microenvironment | Glutamate level regulation, redox balance |
| 10 | *RNLS* | Microenvironment | Regulates cardiac function, blood pressure and immune microenvironment. |
| 10 | *UTF1* | Stem cell | Maintains pluripotency in embryonic stem cells [2] |
| 11 | *DAGLA* | Microenvironment | Affects hormone and immune microenvironment in the CNS |
| 11 | *MAP4K2* | Stem cell | Member of a pathway that affects cell fate of pluripotent stem cells |

| | | | |
|---|---|---|---|
| 11 | *MUC6* | Microenvironment | Forms gelatinous barrier to protect epithelial cells, alters cell signaling, modifies ECM and immunity |
| 14 | *GPX2* | Microenvironment | Redox balance and detoxification |
| 15 | *ANPEP* | Microenvironment | Endocytosis, amino acid and peptide transport. Used as a stem cell marker |
| 16 | *MT3* | Microenvironment | Detoxification, cell response to stress, zinc homeostasis. May be involved in deciding cell fate. |
| 16 | *POLR3K* | Other | Transcription, Subunit in RNA polymerase III. |
| 17 | *CLDN7* | Microenvironment | Mitochondrial tight junctions |
| 17 | *RAC3* | Stem cell | Required to maintain pluripotency of normal stem cells [3] |
| 17 | *RTN4RL1* | Microenvironment | Signaling cascades that impact actin cytoskeleton. May have stem cell like properties in nervous system but too little information available |
| 19 | *EPOR* | Stem cell | Stem cell differentiation especially in erythrocyte and neural progenitor cells [4]. |
| 19 | *GAMT* | Other | Creatine synthesis, muscle function, energy metabolism |
| 19 | *ZNF576* | Other | Transcription regulation |
| 22 | *A4GALT* | Microenvironment | Production of a glycolipid called globotriaosylceramide Membrane composition and receptor availability for cell interactions. |
| 22 | *MAPK8IP2* | Stem cell | Affects cell fate decisions in pluripotent neural progenitor cells, probably affects JNK signal pathway [5] |
| 22 | *RASD2* | Stem cell | Stem cell differentiation pathways |
| 22 | *SOX10* | Stem cell | Stem cell regulator and marker, mammary stem cells [6] |

References

1. Rieskamp JD, Rosado-Burgos I, Christofi JE, Ansar E, Einstein D, Walters AE, et al. Excitatory amino acid transporter 1 supports adult hippocampal neural stem cell self-

renewal. iScience. 2023 Jul 21;26(7):107068. PMID: 37534178. doi: 10.1016/j.isci.2023.107068.
2. Raina K, Dey C, Thool M, Sudhagar S, Thummer RP. An Insight into the Role of UTF1 in Development, Stem Cells, and Cancer. Stem Cell Rev Rep. 2021 Aug;17(4):1280–93. PMID: 33517544. doi: 10.1007/s12015-021-10127-9.
3. Panelo LC, Machado MS, Rubio MF, Jaworski F, Alvarado CV, Paz LA, et al. High RAC3 expression levels are required for induction and maintaining of cancer cell stemness. Oncotarget. 2018 Jan 19;9(5):5848–60. PMID: 29464039. doi: 10.18632/oncotarget.23635.
4. Niemann T, Greiner JFW, Kaltschmidt C, Kaltschmidt B. EPO regulates neuronal differentiation of adult human neural-crest derived stem cells in a sex-specific manner. BMC Neurosci. 2023 Mar 6;24(1):19. PMID: 36879191. doi: 10.1186/s12868-023-00789-1.
5. Roessler R, Goldmann J, Shivalila C, Jaenisch R. JIP2 haploinsufficiency contributes to neurodevelopmental abnormalities in human pluripotent stem cell-derived neural progenitors and cortical neurons. Life Sci Alliance. 2018 Aug;1(4):e201800094. PMID: 30456368. doi: 10.26508/lsa.201800094.
6. Dravis C, Spike BT, Harrell JC, Johns C, Trejo CL, Southard-Smith EM, et al. Sox10 Regulates Stem/Progenitor and Mesenchymal Cell States in Mammary Epithelial Cells. Cell reports. 2015 Sep 29;12(12):2035–48. PMID: 26365194. doi: 10.1016/j.celrep.2015.08.040.
