## Supplemental Table S1 for "Epstein-Barr virus induced epigenetic reprogramming drives cancer stem cell emergence in breast cancer"

***Supplementary Table S1. Matching abnormal methylation of gene controls in breast cancers and EBV cancers affects genes associated with stem cells and the microenvironment. S=Stem cell related, M=microenvironment, Other=not directly related to stem cells or microenvironment.***

| **Chrom-osome** | **CLASS** | **Gene with abnormal methylation in breast cancer corresponding to EBV cancers** | **Function related to stem cell self renewal, differentiation, microenvironment, or other** |
|---|---|---|---|
| 1 | ***S*** | ***EDARADD*** | Guides development of structures that originate from ectodermal stem cells. Critical for interactions between ectoderm and mesoderm during development |
| 1 | ***S*** | ***ETNK2*** | Essential for stem cell pluripotency Synthesis of critical membrane component of both mitochondrial and plasma cell membranes |
| 1 | ***S*** | ***HES4*** | Regulates development especially in the immune system, bone and retina [1]. Responds to NOTCH signaling. Regulated by Wnt and Hedgehog signaling. |
| 1 | ***S*** | ***IGSF21*** | Expressed in stem cell lines. Regulates inhibitory synapse formation in brain [2] |
| 1 | ***S*** | ***KIAA1522*** | Stem cell pluripotency [3] |
| 1 | ***S*** | ***LRRN2*** | Implicated in regulating pluripotency and differentiation [4, 5]. Cell adhesion and signal transduction |
| 1 | ***S*** | ***MACF1*** | Required for beta catenin signaling in nucleus and important for embryonic development. Binds to growing microtubule ends in cytoskeleton, |
| 1 | ***S*** | ***PAQR7*** | Membrane progesterone receptor linked to stem cell ability to regenerate nerve cells [6] |
| 1 | ***S*** | ***SPOCD1*** | Human stem cell fate determination [7]. Directs piRNAs mediated DNA methylation [8] |
| 1 | ***S*** | ***TCEA3*** | Regulates stem cell differentiation potential. Highly expressed in embryonic stem cells [9]. Enhances skeletal muscle differentiation |
| 1 | ***S*** | ***WLS*** | Critical mediator in Wnt pathway insuring their correct trafficking and secretion. Essential for pattering and body axis development |
| 1 | ***M*** | ***ATPAF1*** | Assembly factor for mitochondrial ATP synthase complex in electron transport chain |
| 1 | ***M*** | ***CAMK2N1*** | Mitochondrial stress response [10] |
| 1 | ***M*** | ***C1orf64*** | Androgen receptor target |
| 1 | ***M*** | ***CDC42BPA*** | a kinase regulated by the small GTPase CDC42, involved in actin–myosin cytoskeleton dynamics, which are crucial for cell shape, migration, and adhesion |
| 1 | ***M*** | ***KCNK2*** | Regulates membrane potential found in stem cells. Important for stem cell function but not considered a stem cell gene |
| 1 | ***M*** | ***MRPS21*** | Mitochondrial ribosomal protein |
| 1 | ***M*** | ***PKP1*** | Cell adhesion and communication |
| 1 | ***M*** | ***RGS7*** | Accelerates G protein inactivation, dampening GPCR signaling which mediate responses to extracellular signals. In this way, RGS7 influences cellular perception and response to environmental stimuli. |
| 1 | ***M*** | ***SLC2A1*** | Glucose transporter. One of the transporters that can contribute to the Warburg effect |
| 1 | ***M*** | ***ST6GALNAC5*** | Modifies cell surface interactions i.e. cell-cell, cell-extracellular matrix interactions. Sialyltransferase. Mediates breast cancer brain metastasis [11] |
| 1 | ***M*** | ***TMEM61*** | Regulates actin filopodia formation as in primary cilia, determining how cells move, adhere, and interact with surrounding tissues. |

| | | | |
|---|---|---|---|
| 1 | *Other* | *IL12RB2* | Crucial to activate T-helper cells after they are already committed . Supports stem cell self renewal [12] |
| 2 | ***S*** | ***DNMT3A*** | Stem cell activation and function, found in hematopoietic and embryonic stem cells. Enzyme for DNA methylation of host genes. Critical to control T-cells and regulated by EBV in gastric carcinoma [13, 14] |
| 2 | ***S*** | ***EFHD1*** | Regulates Hippo pathway, which controls stem cell properties . May contribute to stem cell development [15]. Mitochondrial calcium sensor, membrane trafficking. Immune cell activation |
| 2 | ***S*** | ***EN1*** | Homeobox gene. Regulates stem cells, cell growth and differentiation. Controls pattern formation in nervous system development. |
| 2 | ***S*** | ***FHL2*** | Stem cell regulator including hematopoietic and mesenchymal cells. Strongest developmental pathway relationship is to Wnt signaling [16]Essential to maintain self renewal during replication stress. Regulates ovarian cancer tumor progression Weakens gastrointestinal tumor expression [17, 18]. |
| 2 | ***S*** | ***IGFBP5*** | Regulates stem cell differentiation and proliferation |
| 2 | ***S*** | ***LRP1B*** | Regulates neural stem cells. Frequently mutated in cancer by epigenetic or genetic mechanism. Putative tumor suppressor [19]. |
| 2 | ***S*** | ***LYPD6B*** | Expressed in stem cells. Hypermethylation associated with invasive phenotype in melanoma [20] |
| 2 | ***S*** | ***LYPD6*** | Enhances Wnt beta catenin signals [21]interacts with LRP6 a Wnt coreceptor |
| 2 | *S* | ***OTX1*** | Stem cell development |
| 2 | **S** | ***PRKCE*** | Regulates stem cells |
| 2 | ***S*** | ***SH3BP4*** | Stem cell regulation |
| 2 | ***S*** | ***SIX2*** | Direct downstream target of HOXa2 [22] Regulates stem cells |
| 2 | ***S*** | ***SIX3*** | Homeobox gene. Essential for neuronal differentiation, regulates Wnt pathway [23]Cancer suppressor in most cases but can also be a promoter depending on whether pathways inhibit or promote cancer [24] |
| 2 | ***S*** | ***SP5*** | Maintains pluripotent stem cells |
| 2 | ***S*** | ***TCF7L1*** | Regulates pluripotency and lineage specification [25]Embryonic stem cell signature gene |
| 2 | ***S*** | ***VAX2*** | Homeobox gene, controls basic eye development processes. |
| 2 | ***M*** | ***CLEC4F*** | Binds galactose and N-acetyl galactosamine which allows Kupffer cells to recognize and respond to these products from damaged cells or pathogens. Marker for macrophages recruited to mitochondrial damage signals [26].Destroys platelets that have been desialylated by bacterial infection [27]. |
| 2 | ***M*** | ***GAD1*** | Response to inhibiting mitochondrial oxidative phosphorylation [28]. Produces inhibitory neurotransmitter gamma amino butyric acid from glutamic acid. Overexpression a poor prognostic indicator in NPC [29] |
| 2 | ***M*** | ***GPC1*** | Regulates signaling pathways that influence stem cell behavior, including proliferation, differentiation, and migration Involved in EBV infection and immune cell infiltration [30] |
| 2 | ***M*** | ***INHBB*** | Shapes cancer microenvironment [31]. Regulates cell cycle, apoptosis and synthesis of estradiol and progesterone. Correlates with immune infiltration in gastric cancer [32] |
| 2 | ***M*** | ***MBOAT2*** | Shapes membrane fluidity and composition, interactions with ECM. Remodels lecithins, Activates quiescent neural stem cells [33] |
| 2 | ***M*** | ***MYO1B*** | Actin binding motor protein that couples the actin cortex to the plasma membrane, enabling cells to change shape and form protrusions like filopodia. These protrusions are essential for cell migration, especially during development or wound healing. |
| 2 | ***M*** | ***NPAS2*** | Controls glucose metabolism Upregulates glycolytic genes [34]. Circadian clock gene |

| | | | |
|---|---|---|---|
| 2 | *M* | *PXDN* | Extracellular matrix and basement membrane formation. Mammary gland development [35] |
| 2 | *M* | *RAB11FIP5* | Directs proteins to specific locations in the cell surface, influencing how cells respond to external signals. In salivary epithelial cells, RAB11FIP5 mediates the V-ATPase transport to plasma membrane so that cells can respond to acidic conditions.<br>Essential for calcium-stimulated exocytosis in neuroendocrine cells, affecting release of signaling molecules into surroundings. Also affects mitochondrial position and shape [36]NK cell function |
| 2 | *M* | *RAMP1* | Partners with receptors to influence cell response to their surroundings Stress stimulated hematopoiesis [37] |
| 2 | *M* | *SLC16A14* | Orphan gene but close relatives are Monocarboxylate transporter that maintains stem cell niche microenvironment, Pyruvate lactate axis, energy metabolism, and gluconeogenesis |
| 2 | *M* | *SLC1A4* | Transporter for neutral amino acids Ala, Ser, Cys, Thr, Pro, and OH-Pro |
| 2 | *Other* | *DDX11L2* | Originally classified as a pseudogene but may contain a functional promoter. |
| 2 | *Other* | *LONRF2* | Protein quality control, Stem cell quality |
| 2 | *Other* | *MLPH* | Transports intracellular organelles. Facilitates movement along actin filaments. Development of normal pigmentation in skin, hair, eyes. Indirect and context dependent influence on cell microenvironment. |
| 3 | *S* | *PDZRN3* | Development and stability of vascular plexuses, Wnt planar cell polarity [38]. Negative regulator of Wnt pathway [39]. Ubiquitin ligase that marks proteins for destruction. |
| 3 | *S* | *RPL39L* | Ribosomal protein paralog highly expressed in pluripotent cells supports pluripotency and differentiation [40], contributes to differentiation in multiple lineages |
| 3 | *S* | *TNIK* | serine/threonine kinase that activates Wnt signaling pathway, TNIK interacts with transcription factors including TCF4, adjusting gene response to Wnt signals [41] |
| 3 | *S* | *ZIC4* | Transcription factor with important role in development [42]. Target of Wnt pathway |
| 3 | *S* | *ZNF502* | Insufficient information but probably a stem cell gene because it belongs to a family with strong stem cell links and is expressed in germ line cells May be involved in stem cell pluripotency affects cell differentiation |
| 3 | *M* | *CNTN3* | Cell-cell interactions to organize the extracellular environment. Cell adhesion and directional guidance during differentiation of neural stem cells. |
| 3 | *M* | *IGF2BP2* | Regulates glycolysis [43] |
| 3 | *M* | *NMNAT3* | NMNAT3 enzyme catalyzes conversion of NMN to $NAD^+$, primarily within mitochondria. This conversion is essential for redox balance, energy metabolism, and sirtuin activation, especially Sirt3. |
| 3 | *M* | *TRH* | Integrator of energy metabolism [44, 45]. Transcription factor produced in the hypothalamus |
| 4 | *S* | *FAT1* | Loss promotes cancer initiation stem like cells [46], Essential in development for cell polarity, proliferation and survival. Loss is lethal to mice [47] |
| 4 | *S* | *GRIA2* | Cell communication neuron to neuron signaling during brain development. Shapes neuronal circuits [48]. Also expressed in pituitary. |
| 4 | *S* | *NPNT* | Regulates sox2 expressed in bulge stem cells other stem cell processes [49], |
| 4 | *S* | *PPP2R2C* | Interacts with signaling pathways that direct cell fate [50] Regulatory factor that influences development and stem cell behavior. Encodes PP2A enzyme which affects pluripotency and differentiation. |
| 4 | *S* | *PROM1* | Well-known stem cell marker on several types of stem cells [51] |

| | | | |
|---|---|---|---|
| 4 | S | ZFP42 | Maintains undifferentiated stem cells and regulates their differentiation [52] |
| 4 | M | AFAP1 | Cell adhesion, migration. Actin filaments in cytoskeleton, cell signaling |
| 4 | M | HADH | Encodes part of a protein essential for mitochondrial fatty acid oxidation Hydroxyacyl-CoA hydrogenase |
| 4 | M | LMCH1 | Cell motility, cytoskeleton, and adhesion. Muscle stress fibers |
| 4 | M | SHROOM3 | Regulates cell shape through actin binding. Implicated in neural tube defects |
| 5 | S | CXXC5 | An epigenetic regulator that recruits Tet2 demethylase to regulate interferon responses [53]. Modulates the balance between self-renewal and differentiation to influence cell fate decisions [54] |
| 5 | S | IQGAP2 | May contribute to differentiation of stem cells. Participates in MAPK/ERK and PI3K/AKT/mTOR pathways, which are options for cell fate determination. Upregulated in EBV-infected cells. Knockdown of IQGAP2 affects the clumping of infected cells. Varied functions related to its binding partner. Modulates the cytoskeleton [55]. |
| 5 | S | IRX2 | Methylation determines differentiation pathway of induced pluripotent stem cells [56] |
| 5 | S | IRX4 | Multi-potent progenitor cell marker |
| 5 | S | SPRY4 | Gatekeeper of MAPK/ERK responses, one of the pathways that regulate stem cell fate [57]. SPRY4 activity has been detected in embryonic stem cells |
| 5 | M | EPB41L4A | Interactions between the cytoskeleton and the plasma membrane. Contributes to microenvironmental interactions and to beta catenin signaling in cell fate decisions |
| 5 | M | FAM153A | Influences cell adhesion and migration by affecting cell shape [58] |
| 5 | M | FBN2 | Extracellular matrix formation can influence stem cell differentiation |
| 5 | M | NNT | Mitochondrial redox regulation, antioxidant defense |
| 5 | M | TNIP1 | Inflammation regulator, tissue homeostasis, autoimmunity |
| 5 | Other | ANKHD1 | Cell cycle regulation, proliferation and survival |
| 5 | Other | SNORD123 | Guides methylation of ribosomal RNA, indirectly affects stem cells and microenvironment |
| 5 | Other | SUB1 | Safeguards genome stability. |
| 6 | S | GJA1 | Essential for primed stem cell functions to allow communication between cells [59]. Negative regulator of SOX2 expression [60] |
| 6 | S | PHF10 | Essential for chromatin remodeling and hematopoietic stem cell maintenance [61], PHF10 is a key component of the npBAF chromatin remodeling complex, which is essential for the self-renewal and proliferation of multipotent neural stem cells. |
| 6 | S | RBM24 | Regulates embryonic lineage differentiation and cell homeostasis [62]Related to BMP signaling [63]. Driver of lineage commitment to cardiac cells |
| 6 | S | RPS6KA2 | Enhances stem cell differentiation [64]. Promotes cell growth and differentiation. Downstream effector of MAPK/ERK pathway |
| 6 | S | SPDEF | Influences differentiation and maturation of somatic goblet cells. Regulated by NOTCH signals. Enhances cancer stem cell like properties [65] |
| 6 | S | TFAP2A | Causes embryonic stem cells to differentiate to neural crest cells by combining with OCT4-SOX2 complex [66]. Partners with GATA2/3 and TFAP2C in a transcriptional circuit named the TEtra network. This network represses pluripotency genes like OCT4 to differentiates toward trophectoderm (placental lineage). Inhibiting TFAP2A disrupts this balance, preventing proper lineage commitment [67]. TFAP2A binds to epigenetically inactive placental genes at the onset of differentiation, helping activate them. Without TFAP2A, these genes remain silent, stalling differentiation". |
| 6 | S | TFAP2B | Specifies an embryonic melanocyte stem cell that retains adult potential for multiple fates [68] Suppression of Epithelial-to-mesenchymal transition and Wnt/β-catenin pathways in certain cancers [69] |

| | | | |
|---|---|---|---|
| 6 | *S* | *TRERF1* | Induces preleukemic stem cell expansion [70]. Transcriptional-regulating factor 1 is a zinc-finger transcriptional regulating protein which interacts with CBP/p300 to regulate the human gene CYP11A1. |
| 6 | *M* | *CAP2* | actin cytoskeleton regulation, which affects cell shape, movement, and signaling. Interacts with pathways which control cell adhesion. |
| 6 | *M* | *CNKSR3* | Transepithelial sodium transport [GeneCards] Aldosterone mediated sodium transport |
| 6 | *M* | *PACSIN1* | Endocytosis, receptor trafficking |
| *6* | *M* | *PTPRK* | Protein tyrosine phosphatase, controls adhesion molecules that determine stem cell niche interactions [71]targeting removes stem cell function from colon cancer cells [72] Dephosphorylates beta catenin in Wnt pathway [73]. Promotes differentiation as a key modulator and suppresses proliferation. |
| *6* | *M* | *RSPH9* | Cilia function and motility |
| 6 | *M* | *TUBB2B* | Microtubule structure, controls how the cell interacts with its environment. |
| *6* | *Other* | *MTCH1* | Decides if cell should undergo apoptosis. Guides proteins into place in the membrane. Mitochondrial outer membrane protein |
| 6 | *Other* | *PCMT1* | Protein quality control, cell survival |
| 6 | *Other* | *PRSS16* | Serine protease shapes T-cell abilities to recognize antigens |
| *6* | *Unkn* | FAM50B | Identified in gene signatures in stem cells. Development of germ cells [74]. Regulates immune microenvironment Differential methylation relies on histone modifications in embryonic stem cells. Gene is imprinted in humans and associated with retrotransposon insertion sites [70] |
| *6* | *Unkn* | LOC401242 | Too little information available |
| 7 | *S* | *COBL* | Cell polarity and development |
| 7 | *S* | *EPHA1* | Cancer stem cell marker [75] |
| 7 | *S* | *HOXA10-HOXA9* | Both HOXA10 and HOXA9 activate stem cell specific genes [76]. Transcription read through associates with viral infections [77] |
| 7 | *S* | *LFNG* | Differentiation and cell fate [78] |
| 7 | *S* | *NACAD* | Regulates fate of intestinal stem cells [79]. Controls metabolism that stem cell differentiation pathways depend on. |
| 7 | *S* | *SMO* | Regulates stem cell self renewal and development Part of hedgehog pathway [80] |
| 7 | *M* | *CLDN3* | Controls interact with microenvironment and shapes cell architecture. Contributes to cancer stem cell phenotype and highly expressed in cancer stem cells [81] |
| 7 | *M* | *GSTK1* | Glutathione S-transferase, detoxification enzyme |
| 7 | *M* | *NPTX2* | Anchors AMPA receptors at synapses by binding to the extracellular section. This creates a more structured extracellular matrix around the synapse [82]. |
| 7 | *M* | *NRCAM* | Sculptor that shapes stem cell niche microenvironment. Regulates stem cells in the nervous system [83]. |
| 7 | *M* | *PTPRN2* | Phosphatidylinositol phosphatase that alters phosphoinositide levels in the plasma membrane. These levels affect membrane curvature, vesicle formation, exocytosis, and the actin cytoskeleton. PTPRN2 indirectly releases membrane cofilin which cuts actin filaments. Severing these filaments increases cell relocation, invasion, and adaptation to forces acting on the microenvironment. Vesicle-mediated exocytosis, controls signaling from the extracellular matrix, communication between cells, hormone and neurotransmitter release [84] |
| 7 | *M* | *TMEM176A* | Cellular microenvironment in immune cell contexts |
| 7 | *M* | *VIPR2* | Water and ion flux. |

| | | | |
|---|---|---|---|
| 7 | *Other* | *ADCY1* | Adenyl cyclase, may be important to maintain stem cells |
| 7 | *Other* | *ZC3HC1* | Stabilizes FANCD2, a critical member of the FA-BRCA pathway for DNA repair. |
| 8 | ***S*** | ***ADAM32*** | RT-PCR results showed ADAM32 affects expression of genes associated with stemness [85], has a role in development, membrane protein. |
| 8 | ***S*** | ***FBXO16*** | Marks proteins for destruction by ubiquitylation. Disrupts beta catenin signaling in breast cancer [86] |
| 8 | ***S*** | ***FBXO32*** | Regulates stem cell differentiation and renewal [87] |
| 8 | ***S*** | ***LAPTM4B*** | Associated with stemness especially in populations of cancer cells [88]. This differs from its role in normal cells where LAPTM4B is associated with lysosomes and endosomal traffic. |
| 8 | ***S*** | ***NKX3-1*** | Determines cell fate of pluripotent stem cells in specific contexts i.e. fate-determining factor, steering pluripotent stem cells toward a specialized lineage [89] |
| 8 | ***S*** | ***SDR16C5*** | A short-chain dehydrogenase/reductase (SDR) superfamily member that catalyzes the oxidation of retinol (vitamin A) into retinaldehyde—a key early step in the retinoic acid biosynthesis pathway. This enzymatic role is essential for mediating retinoic acid signaling, which influences numerous developmental and metabolic processes, but SDR16C5 does this as part of a metabolic pathway rather than by acting directly on stem cell properties or by actively remodeling the microenvironment. |
| 8 | ***S*** | ***SQLE*** | Key enzyme in cholesterol biosynthesis. Upregulates markers for stem cell properties [90] |
| 8 | ***M*** | ***ARHGEF10*** | Regulates rho GTPases involved in cell signaling, cytoskeletal dynamics, vesicle trafficking, and neural developmental programs. Interacts with other small GTPases like rad6 and rab8 which control cell shape, polarity and movement [91]. |
| 8 | ***M*** | ***IKBKB*** | Regulates inflammation and immune response activates NF-KB to produce cytokines. |
| 8 | ***M*** | ***LRRC6*** | Development and function of cilia [92] |
| 8 | ***M*** | ***MAL2*** | Modulates membrane traffic to control cell response to microenvironment. Vesicular trafficking, formation of lipid rafts, and signal transduction in T-cells and myelin-producing cells |
| 8 | ***M*** | ***MSRA*** | Protects cells from oxidative stress. A special truncated form protects mouse embryonic stem cells from oxidation [93]. Used as a marker to isolate stem cell populations |
| 8 | ***M*** | ***SLC7A2*** | Regulates the intracellular levels of cationic amino acids such as Arginine, affects cellular metabolism and energy homeostasis. Alterations directly influence the metabolic fitness of nearby immune cells which depend on adequate levels of L-arginine for an immune response. Helps create an immunosuppressive niche environment. Expressed in embryonic and induced pluripotent stem cells. |
| 8 | ***M*** | ***TOP1MT*** | Relieves supercoiling of mitochondrial DNA. Helps tumors adapt to environments with limited resources [94] |
| 8 | *Other* | *FAM110B* | Mitosis control, locates at centrosomes and spindle poles. Might influence stem cell niche remodeling. |
| 8 | *Other* | *RIMS2* | Scaffold protein that regulates release of neurotransmitters and hormones Expressed in stem cells especially during neuronal development [95] Regulates exocytosis. Coordinates vesicle treaffic |
| 8 | *Other* | *TIGD5* | Potential influence on chromatin architecture. DNA transposon expressed in neural stem cells and primordial germ cells. Frequently coamplifies with MYC [96]. May be a tumor suppressor. |
| 8 | *Other* | *ZNF696* | Regulates transcription mediated by RNA polymerase II, Not extensively studied. |
| 9 | ***S*** | ***GNAQ*** | Contributes to an Rspo3/Wnt3a-Lgr4-Gnaq pathway, enhances β-catenin signaling in MLL leukemic stem cells [97] . Also interacts with Hippo pathway |
| 9 | ***S*** | ***MPDZ*** | Inhibits hippo signaling pathway involved in differentiation of stem cells [98] |

| | | | |
|---|---|---|---|
| 9 | *S* | *NFIB* | Regulates stem cell differentiation and activity in stem cell niche [99]. Associated with NFKB1 and important for NFKB signaling in both B-cells and cytotoxic T-cells [100]. |
| 9 | *S* | *NPDC1* | Regulates neuronal cell differentiation and proliferation and influences stem cell behavior |
| 9 | *S* | *OLFM1* | Development and organization of the nervous system, hematopoiesis [101] |
| 9 | *M* | *CARD9* | Bridges pattern receptors and downstream immune signaling, triggers release of inflammatory cytokines. Balances gut microbes and immune responses [102] |
| 9 | *M* | *CRAT* | Balances fat and glucose metabolism and energy homeostasis in mitochondria |
| 9 | *M* | *DNLZ* | Helps cells respond to environmental stresses like elevated temperature. Preserves mitochondrial functions during stress. Chaperone mediated acquisition of protein 3-dimensional structure. |
| 9 | *M* | *ENTPD2* | Architect of the environment around cells. Released in exosomes to hydrolyze extracellular trinucleotides in the external microenvironment. Overexpression prevents myeloid derived monocyte suppressor cells from differentiating [103] Influence stem cell behavior in taste buds [104] |
| 9 | *M* | *FUT7* | Adhesive interactions of bone marrow cells to regulate distribution, growth, and development in hematopoietic system [105]. Can also affect microenvironment by control of leukocyte trafficking under conditions of inflammation |
| 9 | *M* | *GCNT1* | Glycosylation essential in mucins, and selectins, affecting cell communications |
| 9 | *M* | *GNA14* | Shapes the tumor microenvironment [106] |
| 9 | *M* | *LPAR1* | G protein coupled receptor that triggers cell signaling in response to the environment controls cell migration, interactions, survival and differentiation. Enhances differentiation of neural stem cells, links to multiple sclerosis, an EBV mediated disease [107]. Stress response in stem cells, transplantation survival, also affects microenvironment [108]. |
| 9 | *M* | *PIP5K1B* | Actin filament organization determining cell shape, adhesion and mobility. Contributes to endocytosis and exocytosis. Calcium signaling lipid kinase essential for mitochondrial functions as well as cell signaling |
| 9 | *M* | *PKN3* | Rho GTPase effector which regulates cytoskeletal structure involved in cell micgration and interactions [109] |
| 9 | *M* | *SECISBP2* | Control of selenoprotein translation. Essential to synthesize proteins that respond to oxidative stress in the cell microenvironment. |
| 9 | *M* | *SUSD3* | Regulates how cell-cell and cell environment interactions, influences spread and invasion of tumors. Promotes estrogen dependent cell multiplication and cell interaction and migration. Expressed in stem cells |
| 9 | *Other* | *DAPK1* | Regulates autophagy and apoptosis. Affects cel fate decisions under stress. Involved in cancer stem cell formation also affects tumor stromal cells [110] Downregulation promotes stem cell properties of cancer cells Cell death associated [111] |
| 10 | *S* | *DDIT4* | Regulates stem cell fate links HIF and mTOR pathways [112] |
| 10 | *S* | *DOCK1* | "Exclusively abundant in hematopoietic stem cells" [113] |
| 10 | *S* | *DPYSL4* | Regulates morphogenesis in tooth germ cells [114]. Semaphorin stimulates RAS/MEK/ERK pathway |
| 10 | *S* | *DUSP5* | Regulates MAPK signaling to control cell signaling and differentiation, growth and survival. Phosphatase that regulates signaling pathways, promotes osteogenic differentiation of bone marrow stromal cells [115]. Gatekeeper of developmental and immune processes [116] |
| 10 | *S* | *GATA3* | Regulates self-renewal and cell fate in hematopoietic stem cells [117]. Controlled by BMP signaling and Hh pathways in developing neural crest cells in zebrafish [118] |
| 10 | *S* | *GFRA1* | Neuron survival and differentiation, essential for self-renewal [119] |

| | | | |
|---|---|---|---|
| 10 | *S* | *H2AFY2* | Critical for the differentiation of hematopoietic stem cells [120] Epigenetic regulator that modifies chromatin structure. |
| 10 | *Other* | *LOC283070* | Long non-coding RNA involved in converting prostate cancer cells to androgen receptor independence. Might affect pathways in stem cells that do not require androgen. Probably influences chromatin structure |
| 10 | *S* | *PLXDC2* | Stem cell plasticity, Hematopoietic stem cell marker [121] |
| 10 | *S* | *SPAG6* | Affects TGF-beta [122]and PI3K/AKT [123] pathways Hypermethylated during differentiation of human embryonic stem cells [124]. Identified as a breast cancer biomarker [125] |
| 10 | *S* | *TCF7L2* | Part of Wnt signaling pathway to maintain normal somatic stem cell homeostasis, also involved in embryonic development [126, 127] |
| 10 | *S* | *ZMIZ1* | Modulates activity of stem cell pathways through, NOTCH1, SAD3/4, and beta catenin [128]Transcriptional coactivator that is critical for T-cell development |
| 10 | *S* | *ZNF503* | Essential for embryonic development. Activates MYC, negative regulator of GATA3 potential role in breast cancer stem cell like behavior [129] |
| 10 | *M* | *ARHGAP21* | Negative regulator of Rho GTPase mediated adhesion and migration, essential for hematopoietic stem cell microenvironment [130] |
| 10 | *M* | *ASB13* | Ubiquitin ligase that tags proteins for degradation, may mediate cytokine signaling Knockout increases cell travels and disrupts polymerization of F-actin [131]. ASB13 is part of an axis that regulates how cells respond to the microenvironment. |
| 10 | *M* | *CALY* | Calcium signaling a key part of the stem cell environment at various stem cell differentiation stages. |
| 10 | *M* | *KCNMA1* | Affects stem cell behavior by encoding a conductance channel to regulate nerve and muscle functions. Affects cell microenvironment |
| 10 | *M* | *PPP1R3C* | Regulates gluconeogenesis and glycogen synthesis. Part of a phosphatase complex |
| 10 | *M* | *RGS10* | Regulates G-protein signaling |
| 10 | *Other* | *FAM196A / INSYN2A* | May be involved in chromosome segregation and DNA repair. Regulates inhibitory neurotransmission, scaffolding component at synapses. |
| 10 | *Other* | *TACC2* | Organizes microtubules and couples nucleus to centrosome. Androgen responsive cell cycle regulator [132] |
| 10 | *Other* | *THNSL1* | Threonine biosynthesis |
| 10 | *Unkn* | *C10orf67* | Probably encodes a mitochondrial protein of unknown function |
| 11 | *S* | *ASCL2* | Transcription factor that drives cell fate decisions, highly expressed in stem cells and essential to maintain them [133] |
| 11 | *S* | *IFITM1* | Highly expressed in human embryonic stem cells to maintain pluripotency and to protect pluripotent cells from viral infection [134, 135] Encodes protein that restricts cellular entry by diverse viral pathogens [20, 21]. .IFITMs are part of the innate immune system |
| 11 | *S* | *LGR4* | Regulator that affects stem cell lineage decisions and behavior. Receptor that modulates the Wnt/beta catenin pathway. Loss of LGR4 interferes with the ability of mammary stem cells to repopulate by affecting SOX2 [136] |
| 11 | *S* | *MOB2* | A component of the Hippo signaling pathway that plays a crucial role in stem cell self renewal and differentiation [137]. |
| 11 | *S* | *PLEKHB1* | Membrane associated signaling functions that might indirect regulate cell differentiation pathways. Could control differentiation in specific cell types. Other genes in the same pathway have stronger links to differentiation decisions |

| | | | |
|---|---|---|---|
| 11 | *S* | *TP53I11* | Suppresses extracellular matrix independent survival and mesenchymal transitions in mammary epithelial cells |
| 11 | *S* | *WT1* | Regulates differentiation of progenitor cells, highly expressed in stem cells [138] |
| 11 | *M* | *CHST1* | Carbohydrate sialyltransferase that affects the structure of components of the extracellular matrix which is critical to form the stem cell niche. |
| 11 | *M* | *CORO1B* | Cytoskeleton organization, cell motility. Cell structures critical for cell microenvironment, endothelial networks and tissue regeneration [139] |
| 11 | *M* | *EHD1* | Endocytosis |
| 11 | *M* | *IGSF9B* | Cell-cell adhesion, associated with stem cells |
| 11 | *M* | *MYEOV* | Recruitment and behavior of immune infiltration |
| 11 | *M* | *SYT12* | Helps cells respond to environmental stress Helps cells interpret and respond to surroundings. Correlates with immune cell infiltration in gastric cancer [140] |
| 11 | *M* | *SYT9* | Cellular response to environmental signals. Vesicle mediated transport, clathrin endocytosis |
| 11 | *M* | *TSKU* | Key signaling pathways in the cell microenvironment |
| 11 | *Other* | *ABTB2* | Cell response to toxins |
| 11 | *Other* | *ENDOD1* | Nucleic acid processing and genomic integrity |
| 11 | *Other* | *FADS2* | Encodes rate limiting enzyme for biosynthesis of polyunsaturated fatty acids |
| 11 | *Other* | *TNNI2* | Protects telomeres, regulates their length |
| 12 | *S* | *ATF7IP* | Crucial to regulate hematopoietic stem and progenitor cells by interacting with a histone methyltransferase [141] |
| 12 | *S* | *FAIM2* | Contributes to embryonic stem cells differentiating to specific lineage [142] Marker of immune infiltration [143] |
| 12 | *S* | *FBXO21* | Regulates maintenance and differentiation of hematopoietic stem cells [144]. Regulates epithelial mesenchymal transition [145] |
| 12 | *S* | *HOTAIR* | Epigenetic regulator, that can trigger EMT and regulate stemness of cancer cells [146] Master regulator of chromatin dynamics [147], regulates WNT pathway [148] |
| 12 | *S* | *HOXC10* | In mesenchymal stem cells used as a marker, regulator of body plan [149] |
| 12 | *S* | *NAB2* | Female germline stem cells, brain development [150] |
| 12 | *S* | *NTN4* | Promotes and regulates some stem cell populations [151] |
| 12 | *S* | *WNT5B* | Expressed in stem cells and regulates behavior, often linked to cancer stem cells [152] |
| 12 | *S* | *ZCCHC8* | Regulatory factor in embryonic stem cells [153] |
| 12 | *M* | *CPNE8* | Encoded protein binds to cell membrane in response to calcium signals. Enhances fibroblast and immune cell infiltration of gastric tumor environment. |
| 12 | *M* | *FAR2* | Fatty acid metabolism Defects in FAR2 can cause mitochondrial dysfunction [154]. Affects membrane fluidity and interactions with the cell microenvironment |
| 12 | *Other* | *CHFR* | Cell cycle checkpoint. Has some connection to stem cells because of its role in early germ cell maintenance. |
| 12 | *Other* | *RERG* | Cell growth gene regulated by estrogen |
| 13 | *S* | *DACH1* | Regulates self renewal and differentiation [155]. Enhances cell cycle arrest by binding p53. Prevents breast cancer invasion and metastasis. DACH1 is a cytokine hijacked by a prototype foamy virus [156]. |
| 13 | *S* | *FLT3* | Essential for normal stem cell development and immunity, Hematopoiesis [157]. |
| 13 | *S* | *PCID2* | Suppresses developmental genes to maintain pluripotency in embryonic stem cells. Prevents premature differentiation [158]. |

| | | | |
|---|---|---|---|
| 13 | ***S*** | ***TNFSRF19*** | Actively contributes to stem cell maintenance and differentiation Developmental signaling, modulation of apoptosis, and interactions with other signaling molecules such as TRAF family members also called "TROY" gene [159]. |
| 13 | ***S*** | ***ZIC2*** | Required for embryonic stem cell specification [160] |
| 13 | ***M*** | ***FAM155A*** | Direct structural role in ion channels that control how neurons respond to the microenvironment. Expressed in embryonic stem cells, expression altered as cells differentiate. |
| 13 | ***M*** | ***GRTP1*** | Mitochondrial function and interaction, glucose metabolism, cell growth and differentiation. Connected to insul/TOR/S6K pathway |
| 13 | ***M*** | ***MBNL2*** | Shapes the immune microenvironment of tumors [161]. |
| 13 | *Other* | *BIVM* | DNA endonuclease activity, Probably involved in DNA repair |
| 14 | ***S*** | ***BMP4*** | Encodes critical molecule that guides stem cell differentiation commits multi-potent stem cells to specific differentiation pathway [162] |
| 14 | ***S*** | ***C14orf39/S6OS1*** | Development and function of neural stem cells. Spindle checkpoint essential for sperm and ovarian function [163] |
| 14 | ***S*** | ***CRIP1*** | Myometrial stem progenitor cell population [164] |
| 14 | ***S*** | ***CRIP2*** | Stem cell gene that affects vascular development and development of various tissues [165] |
| 14 | ***S*** | ***MTA1*** | Ability of stem cells to propagate [166]. Stem cell fate [167]. Deacetylates histone H3K27 |
| 14 | ***M*** | ***GPR68*** | Receptor that senses extracellular hydrogen ions. |
| 14 | ***M*** | ***SLC7A8*** | Transporter for neutral amino acids, may influence metabolism during differentiation Essential for oxidative phosphorylation, key supplier of amino acids for innate immunity cells [168] |
| 14 | ***M*** | ***TMEM121*** | Trans membrane protein that responds to and regulates cell behavior that affects surrounding environment, especially in cancer. |
| 15 | ***S*** | ***ALDH1A3*** | Essential for retinoic acid production (pathway to guide HOX gene directed differentiation) Cancer stem cell marker |
| 15 | ***S*** | ***CRABP1*** | Growth factor sensitivity and stemness [169] |
| 15 | ***S*** | ***IGF1R*** | Crucial to establish stem cell pluripotency and self-renewal. Modulates key pathways like PI3K/AKT, RAS/MAPK, and JAK/STAT in stem cells. Niche remodeling [170] |
| 15 | ***S*** | ***NR2F2*** | Regulates differentiation and immune control in stem cells [171]. A long non-coding RNA downregulated in EBV infection [172] |
| 15 | ***S*** | ***PCSK6*** | Protease needed to activate stem cell factors BMP and Nodal [173] |
| 15 | ***S*** | ***WDR72*** | Enhances stem cell properties of lung cancer cells. Activates AKT/HIF signaling [174] |
| 15 | ***M*** | ***HDGFRP3*** | Extracellular signaling that remodels the extracellular environment [175] |
| 15 | ***M*** | ***KLF13*** | Releases chemokines that shape the local environment |
| 15 | ***M*** | ***SLCO3A1*** | Moves organic anions across cell membranes. Part of a shared gene signature across multiple stem cell lines [176] |
| 16 | ***S*** | ***FBXL16*** | Embryonic stem cell differentiation [177] |
| 16 | **S** | **IRX3** | Hematopoietic differentiation, Neurogenesis in hypothalamus [178], stem cell behavior |
| 16 | **S** | **IRX5** | Hematopoietic differentiation, Neurogenesis in hypothalamus , stem cell behavior |
| 16 | ***S*** | ***NETO2*** | Significant role in regulating multiple stem cell pathways Wnt, JAK-STAT, MAPK/AKT and TGF-beta [179] although not a core stem cell gene. |
| 16 | ***S*** | ***TOX3*** | Regulates neural stem cell identity [180] |
| 16 | ***M*** | ***C16orf45*** | Cell migration, cell shape, protein-protein interactions |

| | | | |
|---|---|---|---|
| 16 | *M* | *CLEC16A* | Autophagy, mitophagy, regulates normal glucose responsive insulin release, inflammation, endosomal trafficking |
| 16 | *M* | *MSLN* | Membrane antigen, cell adhesion and recognition. |
| 16 | *M* | *RAB40C* | GTPase involved in protein degradation**,** membrane trafficking**, and** cell migration |
| 16 | M | *VPS35* | Sorting and trafficking proteins, essential for mitochondrial function and fusion |
| 16 | M | *WDR90* | Centrosome and cilium structure sensing microenvironment |
| 16 | Other | *APRT* | Purine salvage pathway |
| 16 | Other | *RHBDL1* | Protein degradation and signaling |
| 16 | *Other* | *Z3H18* | RNA processing and turnover |
| 17 | *S* | *ANAPC11* | Key regulator of adult keratinocyte stem cell fate and keratinocyte homeostasis [181]. Important in early stage development. Linked to endoderm stem cells Crucial to maintain hematopoietic stem cells. Essential for cell division |
| 17 | *S* | *B4GALNT2* | Stem cell regeneration Infection susceptibility [182] |
| 17 | *S* | *CYB5D2* | Enhances repair of damaged heart cells by pluripotent stem cells after infarction [183]. Expressed in neural stem cells and used as a stem cell marker Predicted to be involved in nervous system development. Likely affects multiple differentiation pathways. |
| 17 | *S* | *FASN* | A metabolic gatekeeper that is essential for the activity of multiple stem cell pathways such as Wnt, NOTCH, and EGF [184]. Influences cell fate decisions thorough metabolic reprogramming. Controls adult neural stem cell activity [185]Cell energy production fatty acid synthase |
| 17 | *S* | *GPRC5C* | Regulates stem cell quiescence of hematopoietic stem cells [186] induced by retinoic acid [GeneCards] |
| 17 | *S* | *JMJD6* | Demethylase that influences chromatin structure, hydroxylates lysine on histones and non-histones. Promotes stem cell self renewal and ability to differentiate into multiple cell types [187] |
| 17 | *S* | *KRT19* | Regulates NOTCH signaling pathway and stem cell properties [188]. Balances Wnt NOTCH signaling [189] |
| 17 | *S* | *LGALS3BP* | Regulates positions of neural progenitors [190]Related to EGF signaling [191] |
| | *S* | *NUP85* | Nuclear architecture in stem cells |
| 17 | *S* | *PMP22* | Regulates stem cell property of self renewal in GC. [192] TEAD1 and YAP/TAZ key hippo effector bind to enhancers upstream of PMP22 [193] |
| 17 | *S* | *PPP1R1B* | Stem cell development |
| 17 | *S* | *PSMC31P* | Germline male stem cell development [Gene Cards] |
| 17 | *S* | *RAC3* | Required to maintain pluripotency of normal stem cells [194] |
| 17 | *S* | *RAI1* | Upregulated in response to retinoic acid signaling for differentiation. Brain development, circadian rhythm, |
| 17 | *S* | *SOX9* | Stem cell factor critical in embryonic development. Immune cell infiltration into tumors [195]. |
| 17 | *S* | *SREBF1* | Regulates hematopoietic stem cell fate, function, and survival [196] |
| 17 | *S* | *UTP6* | Hypermethylation associated with stem-cell like properties in colorectal cancer stem cells Secondary control of Wnt pathway via FOXK2 [197]. May be involved in transcriptional regulation of pluripotency regulatory factors Low expression increases stemness. |
| 17 | *M* | *ATP2A3* | Significant role in controlling cellular microenvironment, and calcium homeostasis and cell signaling. Encodes an ATP-ase enzyme for calcium transport and sequestration. Expressed in embryonic stem cells and in hematopoietic and epithelial cell lineages. |
| 17 | *M* | *ATP5H* | Mitochondrial ATP synthase |
| 17 | *M* | *CDC42EP4* | Organizes cell shape, movement, and pseudopodia formation, Interacts with GTPases, senses environment |

| | | | |
|---|---|---|---|
| 17 | *M* | *CORO6* | Actin dynamics, cytoskeleton, cell growth |
| 17 | *M* | *DCXR* | Promotes aerobic glycolysis [198] |
| 17 | *M* | *GCGR* | Glucagon receptor, cell microenvironment, glucose metabolism, stem cell maintenance [199] |
| 17 | *M* | *MRPS7* | Mitochondrial ribosomes |
| 17 | *M* | *RAB34* | Ciliogenesis, sensing microenvironment |
| 17 | *M* | *WNK4* | Adjusts mitochondrial energetics [200]Regulates renal salt handling [201] |
| 17 | *Other* | *AFMID* | Tryptophan metabolism |
| 17 | *Other* | *ALOX12* | Lipoxygenase that regulates platelets, inflammation |
| 17 | *Other* | *CASC3* | Nonsense mediated mRNA removal. May have some developmental functions [202] |
| 17 | *Other* | *GHDC* | Amino acid ligase activity, lactation, Innate immunity |
| 17 | *Other* | *GRIN2C* | NMDA receptor subunit |
| 18 | *S* | *ANKRD12* | Inhibits nuclear receptors via histone deacetylases [GeneCards]. ANKRD12 silencing promotes invasion [203]. |
| 18 | *S* | *CHD7* | Chromatin remodeling Regulates endodermal and mesodermal development, colocalizes with canonical stem cell factors like OCT4 [204] |
| 18 | *S* | *RAB31* | Fine tunes cell fate decision. Interacts with GLI1, an effector in the hedgehog developmental pathway [205]. Regulates glucose transporter [206]. Membrane traffic, exosome biogenesis, found in a variety of stem cells. |
| 18 | *S* | *YES1* | Hippo pathway effector involved in development, growth, repair and homeostasis |
| 18 | *M* | *CYB5A* | Cytochrome related to electron transport chain in mitochondrial energy production |
| 18 | *M* | *DSC2* | Organization of tissues, stability and even cell signaling in the microenvironment |
| 18 | *M* | *DSC3* | Desmosome proteins that control cell-cell adhesion interactions and might affect stem cell niche. Encode a core part of desmosomes that inherently influence the cell microenvironment |
| 18 | *M* | *DTNA* | Cell membrane stability in muscle and neural tissues. Mechanical stability of cell membrane to facilitate interactions with the extracellular matrix |
| 18 | *M* | *GREB1L* | Glycosylation of estrogen receptor, and promotion of immune cell invasion [207, 208]. Intertwined with cell environment regulation |
| 18 | *M* | *ZBTB7C* | Linked to tumor microenvironment [209]. Regulates metabolism and glucose levels |
| 18 | *Other* | *MOCOS* | Purine and aldehyde metabolism indirect connection to energy production. |
| 19 | *S* | *EPS15L1* | Stem cell development and differentiation, endocytosis [210] |
| 19 | *S* | *FUT3* | Fucosyltransferase 3 forms Lewis antigens essential for embryonic development and differentiation pathways. Adds fucose to glycans linked to stem cell characteristics and affects how they differentiate, may enhance cell fate plasticity. Expression reduced in cancer stem cells vs normal stem cells [211]. Fut2 deficits reduce stemness in intestinal stem cells. Lewis antigens serve as binding sites for some pathogens. |
| 19 | *S* | *KLF2* | Regulates embryonic stem cell-specific gene expression [212] |
| 19 | *S* | *LYPD3* | Drives tumor stemness and immune evasion. Maintains tumor stem cell properties Cell adhesion and migration binds to extracellular matrix |
| 19 | *S* | *NFIX* | Regulates production of intermediate progenitor cells from embryonic stem cells in adults regulates the timing of neural differentiation [213] Involved in the oxidative stress response and cell fate decisions [203]. Binds viral and cellular promoters. |
| 19 | *S* | *NR2F6* | Regulates mouse stem cell hematopoiesis and myelopoiesis [214] Crucial role in glucose metabolism and energy production in brown fat. |
| 19 | *S* | *PEG3* | Parental imprinted gene in stem cells in all tissues tested. Stem cell marker [215], Regulates adipogenesis. |
| 19 | *S* | *REEP6* | Photoreceptor development [216] Retinal homeostasis Downstream effector that fine tunes stem cell fate decision. A specialized developmental gene. |

| | | | |
|---|---|---|---|
| 19 | ***S*** | ***ZFP28*** | Maintains pluripotency in embryonic stem cells [217] |
| 19 | ***S*** | ***ZNF256*** | Represses gene expression during development, fine tunes gene activity at specific stages. Cortical specification (Cortecon repository) |
| 19 | ***S*** | ***ZNF329*** | Neural differentiation, cortical specification, deep layers (Cortecon repository) Links to developmental disorders that are not well-documented [MayMyGenome.com]. |
| 19 | ***S*** | ***ZNF350*** | Recruits KAP1 a corepressor that modifies chromatin to determine stem cell identity and lineage [218]. |
| 19 | **S** | **ZNF415** | Contains a KRAB A box, gene is expressed in stem cell niches, |
| 19 | ***S*** | ***ZNF419*** | Related to tumor stemness according to methylation and characteristics of mRBA expression [219] |
| 19 | ***S*** | ***ZNF439*** | Neural differentiation, cortical specification, deep layers (Cortecon repository) |
| 19 | ***S*** | ***ZNF541*** | Meiosis regulator in germ line male cells, likely stem cell gene for sperm cell fate |
| 19 | ***S*** | ***ZNF549*** | Expressed in primordial germ cells, cortical plate, suggesting a role in stem cell differentiation |
| 19 | ***S*** | ***ZNF570*** | Transcription regulation, potential impact on stem cells but inadequate information |
| 19 | ***S*** | ***ZNF626*** | Neural differentiation, cortical specification, upper layers (Cortecon repository) |
| 19 | ***S*** | ***ZNF880*** | Neural differentiation, cortical specification, upper layers (Cortecon repository) |
| 19 | ***M*** | ***CACNG6*** | Transmembrane regulatory receptor voltage dependent calcium channel. Ion homeostasis. |
| 19 | ***M*** | ***CCDC61*** | In epithelial cells with multi cilia, helps maintain interface between cell and its environment. Centrosome location and assembly [220], essential for cell division Proper construction of basal bodies |
| 19 | ***M*** | ***CHST8*** | Affects how cells interact with their surroundings, hormone and immune responses by catalyzing the sulfation of N-acetyl galactosamine on glycoproteins and glycolipids. Neural stem cell proliferation (results from CRISPR screens). Activated by TGF beta [221]. Loss weakens breast cancer tumor growth [222] |
| 19 | ***M*** | ***FXYD5*** | Subunit of sodium-potassium pump, ion homeostasis an membrane potential Confers properties of stem cells [223] |
| 19 | ***M*** | ***GNA11*** | Signal transduction and calcium regulation. Melanocyte development and melanoma [224] |
| 19 | ***M*** | ***KCNK6*** | Potassium channel |
| 19 | ***M*** | ***MYH14*** | Cytokinesis, cell motility and cell polarity. Interacts with ac tin filaments to control how cells respond to their microenvironment |
| 19 | ***M*** | ***PCSK4*** | Helps specify early development environment by processing growth factors and hormones. Specifies cell fates in placental development |
| 19 | *Other* | *KLK5* | Skin maintenance after terminal differentiation of keratinocytes. Keratinocyte differentiation and desquamation |
| 19 | *Other* | *LOC113230* | Non-coding RNA that affects transcription and gene regulation Inhibits arginine synthesis, controlled by TGF-beta induction |
| 19 | ***Other*** | ***MIA-RAB4B*** | Sequesters miRNA as a miRNA sponge to regulate gene expression |
| 19 | ***Other*** | *REXO1* | RNA processing |
| 19 | *Other* | *SSBP4* | Single strand DNA binding, DNA repair |
| 19 | Unkn | LOC100505715 | unknown |
| 20 | ***S*** | ***BMP2*** | Guides stem cell differentiation fate, "shaping embryonic development, maintaining tissue homeostasis, and influencing disease progression" [225]. |
| 20 | ***S*** | ***CEBPB*** | Maintains stem cell self renewal and quiescence, regulates genes associated with cell cycle arrest [226]. |

| | | | |
|---|---|---|---|
| 20 | ***S*** | ***EYA2*** | Self-renewal [227] |
| 20 | ***S*** | ***PTPRT*** | Hematopoietic stem cell self renewal [228] |
| 20 | ***M*** | ***COMMD7*** | Modulates NF-KB pathway for inflammation. Maintains cell surface levels of receptors, directly affecting cell responses to the microenvironment. |
| 20 | ***M*** | ***GPCPD1*** | Controls metabolism to influence cell microenvironment Glycerophospholipid metabolism |
| 20 | ***M*** | ***MAP1L3CA*** | Microtubule interactions, autophagy |
| 20 | ***M*** | ***NECAB3*** | Regulates beta amyloid production. Interacts with dopamine receptor Activates HIF to use glycolysis at normal levels of oxygen [229] |
| 20 | ***M*** | ***SLC24A3*** | Listed as a cancer stem cell gene [230]. Calcium and sodium balance inside cells exchanges with extracellular environment |
| 20 | ***M*** | ***SRXN1*** | Oxidative stress response |
| 20 | *Other* | *BCAS4* | Associated with intermediate filaments |
| 20 | *Other* | *FRG1B* | Pseudogene that may play a role in chromatin remodeling, Elements near FRG1B bind to OCT4 and NANOG. [GeneCards] |
| 20 | *Other* | *RIMS4* | Neuronal activity and synaptic regulation |
| 21 | ***S*** | ***RIPK4*** | RIPK4 activates IRF6, to downregulate proliferative signals (e.g. p63) and promote keratinocyte differentiation. RIPK4 promotes epidermal differentiation, initiating hippo signaling cascade by phosphorylating LATS1/2, inhibiting YAP/TAZ. RIPK4 also enhances differentiation signaling, recruiting LATS1/2 into liquid condensates [231, 232]. |
| 21 | ***M*** | ***ABCG1*** | Mitochondrial function, lipid homeostasis, Removes excess cholesterol from tissues influences stem cell behavior |
| 21 | ***M*** | ***SLC37A1*** | Phosphate regulator, glucose homeostasis |
| 21 | ***M*** | ***TFF1*** | Secreted by gastric mucous cells to stabilize and protect the mucus layer and protect epithelial surfaces. |
| 21 | ***M*** | ***TFF3*** | Mitochondrial function. Protects intestinal barrier |
| 21 | *Other* | *CSTB* | Inhibits cathepsins to regulate protein turnover within cells |
| 22 | ***S*** | ***CCDC117*** | Stem cell fate determination, cardiac morphogenesis [233, 234] |
| 22 | ***S*** | ***FAM83A*** | Associated with stem cell traits in cancer, activates stem cell pathways such as Wnt-beta-catenin and TGF-beta. |
| 22 | ***S*** | ***TBX1*** | Regulates stem cell quiescence vs. proliferation [235], acts by controlling the stem cell gene SOX2. |
| 22 | ***M*** | ***CACNA11*** | Calcium flux |
| 22 | ***M*** | ***CDC42EP1*** | Effector for CDC42 which controls the actin cytoskeleton |
| 22 | ***M*** | ***EMID1*** | Modifies extracellular environment, extracellular matrix interactions |
| 22 | ***M*** | ***KCTD17*** | Formation of primary cilia on cells which sense and respond to the microenvironment |
| 22 | ***M*** | *PANX2* | Regulates cellular processes within the stem cell niche, including cell communication and signaling pathways. Facilitate release of extracellular messengers to modulate extracellular environment |
| 22 | ***M*** | *RAB36* | A GTPase required for stem cell maintenance and cell migration in epithelium of digestive tract [236] |
| 22 | *Other* | *CARD10* | Modulates cytokine and chemokine expression |
| 22 | *Other* | *DMC1* | DNA meiotic recombinase gene expressed in cells undergoing meiosis, including stem cells |
| 22 | *Other* | *XBP1* | Cell stress response, misfolded and unfolded protein metabolism, protein quality control. Under some conditions SBP1 can influence stem cell behavior |

28. Myers MA, Georgiou HM, Byron S, Esposti MD. Inhibition of mitochondrial oxidative phosphorylation induces hyper-expression of glutamic acid decarboxylase in pancreatic islet cells. Autoimmunity. 1999;30(1):43–51. PMID: 10433094. doi: 10.3109/08916939908994759.
29. Lee YY, Chao TB, Sheu MJ, Tian YF, Chen TJ, Lee SW, et al. Glutamate Decarboxylase 1 Overexpression as a Poor Prognostic Factor in Patients with Nasopharyngeal Carcinoma. J Cancer. 2016;7(12):1716–23. PMID: 27698909. doi: 10.7150/jca.15667.
30. Xiong J, Dai YT, Wang WF, Zhang H, Wang CF, Yin T, et al. GPCR signaling contributes to immune characteristics of microenvironment and process of EBV-induced lymphomagenesis. Sci Bull (Beijing). 2023 Nov 15;68(21):2607–19. PMID: 37798178. doi: 10.1016/j.scib.2023.09.029.
31. Jin Y, Cai Q, Wang L, Ji J, Sun Y, Jiang J, et al. Paracrine activin B-NF-kappaB signaling shapes an inflammatory tumor microenvironment in gastric cancer via fibroblast reprogramming. J Exp Clin Cancer Res. 2023 Oct 19;42(1):269. PMID: 37858201. doi: 10.1186/s13046-023-02861-4.
32. Yu W, He G, Zhang W, Ye Z, Zhong Z, Huang S. INHBB is a novel prognostic biomarker and correlated with immune infiltrates in gastric cancer. Front Genet. 2022;13:933862. PMID: 36118865. doi: 10.3389/fgene.2022.933862.
33. Xiaoai Zhao XY, Kévin Contrepois, Francesco Vallania, Mathew Ellenberger, Chloe M. Kashiwagi, Stephanie D. Gagnon, Cynthia J. Siebrand, Matias Cabruja, Gavin M. Traber, Andrew McKay, Daniel Hornburg, Purvesh Khatri, Michael P. Snyder, Richard N. Zare, Anne Brunet. Lipidomic profiling reveals age-dependent changes in complex plasma membrane lipids that regulate neural stem cell aging. Biorxiv. 2022.
34. Bao X, Zhang J, Huang G, Yan J, Xu C, Dou Z, et al. The crosstalk between HIFs and mitochondrial dysfunctions in cancer development. Cell Death Dis. 2021 Feb 26;12(2):215. PMID: 33637686. doi: 10.1038/s41419-021-03505-1.
35. Hao YY, Xiao WQ, Zhang HN, Yu NN, Park G, Han YH, et al. Peroxiredoxin 1 modulates oxidative stress resistance and cell apoptosis through stemness in liver cancer under non-thermal plasma treatment. Biochem Biophys Res Commun. 2024 Dec 17;738:150522. PMID: 39154551. doi: 10.1016/j.bbrc.2024.150522.
36. Landry MC, Champagne C, Boulanger MC, Jette A, Fuchs M, Dziengelewski C, et al. A functional interplay between the small GTPase Rab11a and mitochondria-shaping proteins regulates mitochondrial positioning and polarization of the actin cytoskeleton downstream of Src family kinases. J Biol Chem. 2014 Jan 24;289(4):2230–49. PMID: 24302731. doi: 10.1074/jbc.M113.516351.
37. Suekane A, Saito Y, Nakahata S, Ichikawa T, Ogoh H, Tsujikawa K, et al. CGRP-CRLR/RAMP1 signal is important for stress-induced hematopoiesis. Sci Rep. 2019 Jan 23;9(1):429. PMID: 30674976. doi: 10.1038/s41598-018-36796-0.
38. Sewduth RN, Jaspard-Vinassa B, Peghaire C, Guillabert A, Franzl N, Larrieu-Lahargue F, et al. The ubiquitin ligase PDZRN3 is required for vascular morphogenesis through Wnt/planar cell polarity signalling. Nat Commun. 2014 Sep 8;5:4832. PMID: 25198863. doi: 10.1038/ncomms5832.
39. Konopelski Snavely SE, Susman MW, Kunz RC, Tan J, Srinivasan S, Cohen MD, et al. Proteomic analysis identifies the E3 ubiquitin ligase Pdzrn3 as a regulatory target of Wnt5a-Ror signaling. Proc Natl Acad Sci U S A. 2021 Jun 22;118(25). PMID: 34135125. doi: 10.1073/pnas.2104944118.
40. Wong QW, Li J, Ng SR, Lim SG, Yang H, Vardy LA. RPL39L is an example of a recently evolved ribosomal protein paralog that shows highly specific tissue expression patterns and is upregulated in ESCs and HCC tumors. RNA Biol. 2014;11(1):33–41. PMID: 24452241. doi: 10.4161/rna.27427.

131. Fan H, Wang X, Li W, Shen M, Wei Y, Zheng H, et al. ASB13 inhibits breast cancer metastasis through promoting SNAI2 degradation and relieving its transcriptional repression of YAP. Genes & development. 2020 Oct 1;34(19-20):1359–72. PMID: 32943576. doi: 10.1101/gad.339796.120.
132. Takayama K, Horie-Inoue K, Suzuki T, Urano T, Ikeda K, Fujimura T, et al. TACC2 is an androgen-responsive cell cycle regulator promoting androgen-mediated and castration-resistant growth of prostate cancer. Mol Endocrinol. 2012 May;26(5):748–61. PMID: 22456197. doi: 10.1210/me.2011-1242.
133. van der Flier LG, van Gijn ME, Hatzis P, Kujala P, Haegebarth A, Stange DE, et al. Transcription factor achaete scute-like 2 controls intestinal stem cell fate. Cell. 2009 Mar 6;136(5):903–12. PMID: 19269367. doi: 10.1016/j.cell.2009.01.031.
134. Fu Y, Zhou Z, Wang H, Gong P, Guo R, Wang J, et al. IFITM1 suppresses expression of human endogenous retroviruses in human embryonic stem cells. FEBS Open Bio. 2017 Aug;7(8):1102–10. PMID: 28781951. doi: 10.1002/2211-5463.12246.
135. Friedlova N, Zavadil Kokas F, Hupp TR, Vojtesek B, Nekulova M. IFITM protein regulation and functions: Far beyond the fight against viruses. Front Immunol. 2022;13:1042368. PMID: 36466909. doi: 10.3389/fimmu.2022.1042368.
136. Wang Y, Dong J, Li D, Lai L, Siwko S, Li Y, et al. Lgr4 regulates mammary gland development and stem cell activity through the pluripotency transcription factor Sox2. Stem Cells. 2013 Sep;31(9):1921–31. PMID: 23712846. doi: 10.1002/stem.1438.
137. Delgado ILS, Carmona B, Nolasco S, Santos D, Leitao A, Soares H. MOB: Pivotal Conserved Proteins in Cytokinesis, Cell Architecture and Tissue Homeostasis. Biology (Basel). 2020 Nov 24;9(12). PMID: 33255245. doi: 10.3390/biology9120413.
138. Martinez-Estrada OM, Lettice LA, Essafi A, Guadix JA, Slight J, Velecela V, et al. Wt1 is required for cardiovascular progenitor cell formation through transcriptional control of Snail and E-cadherin. Nat Genet. 2010 Jan;42(1):89–93. PMID: 20023660. doi: 10.1038/ng.494.
139. Werner AC, Weckbach LT, Salvermoser M, Pitter B, Cao J, Maier-Begandt D, et al. Coronin 1B Controls Endothelial Actin Dynamics at Cell-Cell Junctions and Is Required for Endothelial Network Assembly. Front Cell Dev Biol. 2020;8:708. PMID: 32850828. doi: 10.3389/fcell.2020.00708.
140. Niu X, Ma F, Li F, Wei C, Zhang J, Gao Z, et al. Integration of bioinformatics and cellular experiments unveils the role of SYT12 in gastric cancer. BMC cancer. 2024 Oct 29;24(1):1331. PMID: 39472897. doi: 10.1186/s12885-024-13077-w.
141. Wu J, Li J, Chen K, Liu G, Zhou Y, Chen W, et al. Atf7ip and Setdb1 interaction orchestrates the hematopoietic stem and progenitor cell state with diverse lineage differentiation. Proc Natl Acad Sci U S A. 2023 Jan 3;120(1):e2209062120. PMID: 36577070. doi: 10.1073/pnas.2209062120.
142. Littleton SH, Trang KB, Volpe CM, Cook K, DeBruyne N, Maguire JA, et al. Variant-to-function analysis of the childhood obesity chr12q13 locus implicates rs7132908 as a causal variant within the 3' UTR of FAIM2. Cell Genom. 2024 May 8;4(5):100556. PMID: 38697123. doi: 10.1016/j.xgen.2024.100556.
143. Cai J, Ye Z, Hu Y, Wang Y, Ye L, Gao L, et al. FAIM2 is a potential pan-cancer biomarker for prognosis and immune infiltration. Front Oncol. 2022;12:998336. PMID: 36185230. doi: 10.3389/fonc.2022.998336.
144. Dobish KK, Wittorf KJ, Swenson SA, Bean DC, Gavile CM, Woods NT, et al. FBXO21 mediated degradation of p85alpha regulates proliferation and survival of acute myeloid leukemia. Leukemia. 2023 Nov;37(11):2197–208. PMID: 37689825. doi: 10.1038/s41375-023-02020-w.
145. Yang W, Jing T, Wu C, Lu M, Yao X, Xia D, et al. F-box protein FBXO21 overexpression inhibits the proliferation and metastasis of clear cell renal cell carcinoma and is closely related to

the CREB pathway and tumor immune cell infiltration. J Transl Med. 2025 Mar 15;23(1):335. PMID: 40089779. doi: 10.1186/s12967-025-06356-y.
146. Hajjari M, Salavaty A. HOTAIR: an oncogenic long non-coding RNA in different cancers. Cancer Biol Med. 2015 Mar;12(1):1–9. PMID: 25859406. doi: 10.7497/j.issn.2095-3941.2015.0006.
147. Bhan A, Mandal SS. LncRNA HOTAIR: A master regulator of chromatin dynamics and cancer. Biochim Biophys Acta. 2015 Aug;1856(1):151–64. PMID: 26208723. doi: 10.1016/j.bbcan.2015.07.001.
148. Hakami MA, Hazazi A, Abdulaziz O, Almasoudi HH, Alhazmi AYM, Alkhalil SS, et al. HOTAIR: A key regulator of the Wnt/beta-catenin signaling cascade in cancer progression and treatment. Pathol Res Pract. 2024 Jan;253:154957. PMID: 38000201. doi: 10.1016/j.prp.2023.154957.
149. Bhatlekar S, Fields JZ, Boman BM. Role of HOX Genes in Stem Cell Differentiation and Cancer. Stem Cells Int. 2018;2018:3569493. PMID: 30154863. doi: 10.1155/2018/3569493.
150. Corgiat EB, List SM, Rounds JC, Corbett AH, Moberg KH. The RNA-binding protein Nab2 regulates the proteome of the developing Drosophila brain. J Biol Chem. 2021 Jul;297(1):100877. PMID: 34139237. doi: 10.1016/j.jbc.2021.100877.
151. Huang YH, Wang WL, Wang PH, Lee HT, Chang WW. EXOSC5 maintains cancer stem cell activity in endometrial cancer by regulating the NTN4/integrin beta1 signalling axis. Int J Biol Sci. 2024;20(1):265–79. PMID: 38164180. doi: 10.7150/ijbs.86275.
152. Suthon S, Perkins RS, Bryja V, Miranda-Carboni GA, Krum SA. WNT5B in Physiology and Disease. Front Cell Dev Biol. 2021;9:667581. PMID: 34017835. doi: 10.3389/fcell.2021.667581.
153. Wu Y, Liu W, Chen J, Liu S, Wang M, Yang L, et al. Nuclear Exosome Targeting Complex Core Factor Zcchc8 Regulates the Degradation of LINE1 RNA in Early Embryos and Embryonic Stem Cells. Cell Rep. 2019 Nov 19;29(8):2461–72 e6. PMID: 31747613. doi: 10.1016/j.celrep.2019.10.055.
154. Kim JA. Peroxisome Metabolism in Cancer. Cells. 2020 Jul 14;9(7). PMID: 32674458. doi: 10.3390/cells9071692.
155. Hu X, Zhang L, Li Y, Ma X, Dai W, Gao X, et al. Organoid modelling identifies that DACH1 functions as a tumour promoter in colorectal cancer by modulating BMP signalling. EBioMedicine. 2020 Jun;56:102800. PMID: 32512510. doi: 10.1016/j.ebiom.2020.102800.
156. Ma Y, Wei J, Song J, Hu Z, Zhang R, Li Z, et al. The DACH1 Gene Transcriptional Activation and Protein Degradation Mediated by Transactivator Tas of Prototype Foamy Virus. Viruses. 2023 Sep 8;15(9). PMID: 37766305. doi: 10.3390/v15091899.
157. Gilliland DG, Griffin JD. The roles of FLT3 in hematopoiesis and leukemia. Blood. 2002 Sep 1;100(5):1532–42. PMID: 12176867. doi: 10.1182/blood-2002-02-0492.
158. Ye B, Dai Z, Liu B, Wang R, Li C, Huang G, et al. Pcid2 inactivates developmental genes in human and mouse embryonic stem cells to sustain their pluripotency by modulation of EID1 stability. Stem Cells. 2014 Mar;32(3):623–35. PMID: 24167073. doi: 10.1002/stem.1580.
159. Kretzschmar K, Boonekamp KE, Bleijs M, Asra P, Koomen M, Chuva de Sousa Lopes SM, et al. Troy/Tnfrsf19 marks epidermal cells that govern interfollicular epidermal renewal and cornification. Stem Cell Reports. 2021 Sep 14;16(9):2379–94. PMID: 34358453. doi: 10.1016/j.stemcr.2021.07.007.
160. Luo Z, Gao X, Lin C, Smith ER, Marshall SA, Swanson SK, et al. Zic2 is an enhancer-binding factor required for embryonic stem cell specification. Molecular cell. 2015 Feb 19;57(4):685–94. PMID: 25699711. doi: 10.1016/j.molcel.2015.01.007.
161. Gabel AM, Crosse EI, Belleville AE, Hogg SJ, McKellar SA, Abdel-Wahab O, et al. Muscleblind-like proteins are novel modulators of the tumor-immune microenvironment. PLoS One. 2025;20(4):e0321148. PMID: 40305509. doi: 10.1371/journal.pone.0321148.

162. Setiawan AM, Kamarudin TA, Abd Ghafar N. The role of BMP4 in adipose-derived stem cell differentiation: A minireview. Front Cell Dev Biol. 2022;10:1045103. PMID: 36340030. doi: 10.3389/fcell.2022.1045103.
163. Fan S, Jiao Y, Khan R, Jiang X, Javed AR, Ali A, et al. Homozygous mutations in C14orf39/SIX6OS1 cause non-obstructive azoospermia and premature ovarian insufficiency in humans. Am J Hum Genet. 2022 Jul 7;109(7):1343. PMID: 35803236. doi: 10.1016/j.ajhg.2022.06.006.
164. Paul EN, Carpenter TJ, Fitch S, Sheridan R, Lau KH, Arora R, et al. Cysteine-rich intestinal protein 1 is a novel surface marker for human myometrial stem/progenitor cells. Commun Biol. 2023 Jul 3;6(1):686. PMID: 37400623. doi: 10.1038/s42003-023-05061-0.
165. Sun X, Zhang R, Lin X, Xu X. Wnt3a regulates the development of cardiac neural crest cells by modulating expression of cysteine-rich intestinal protein 2 in rhombomere 6. Circ Res. 2008 Apr 11;102(7):831–9. PMID: 18292601. doi: 10.1161/CIRCRESAHA.107.166488.
166. Sato Y, Shibata N, Hashimoto C, Agata K. Migratory regulation by MTA homologous genes is essential for the uniform distribution of planarian adult pluripotent stem cells. Dev Growth Differ. 2022 Apr;64(3):150–62. PMID: 35124813. doi: 10.1111/dgd.12773.
167. Zhang J, Wang Y, Zhang J, Wang X, Liu J, Huo M, et al. The feedback loop between MTA1 and MTA3/TRIM21 modulates stemness of breast cancer in response to estrogen. Cell Death Dis. 2024 Aug 17;15(8):597. PMID: 39154024. doi: 10.1038/s41419-024-06942-w.
168. Panda SK, Kim DH, Desai P, Rodrigues PF, Sudan R, Gilfillan S, et al. SLC7A8 is a key amino acids supplier for the metabolic programs that sustain homeostasis and activation of type 2 innate lymphoid cells. Proc Natl Acad Sci U S A. 2022 Nov 16;119(46):e2215528119. PMID: 36343258. doi: 10.1073/pnas.2215528119.
169. Nagpal I, Wei LN. All-trans Retinoic Acid as a Versatile Cytosolic Signal Modulator Mediated by CRABP1. Int J Mol Sci. 2019 Jul 24;20(15). PMID: 31344789. doi: 10.3390/ijms20153610.
170. Chen PC, Kuo YC, Chuong CM, Huang YH. Niche Modulation of IGF-1R Signaling: Its Role in Stem Cell Pluripotency, Cancer Reprogramming, and Therapeutic Applications. Front Cell Dev Biol. 2020;8:625943. PMID: 33511137. doi: 10.3389/fcell.2020.625943.
171. Ma L, Huang M, Liao X, Cai X, Wu Q. NR2F2 Regulates Cell Proliferation and Immunomodulation in Whartons' Jelly Stem Cells. Genes (Basel). 2022 Aug 16;13(8). PMID: 36011369. doi: 10.3390/genes13081458.
172. Zhang J, Li X, Hu J, Cao P, Yan Q, Zhang S, et al. Long noncoding RNAs involvement in Epstein-Barr virus infection and tumorigenesis. Virology journal. 2020 Apr 9;17(1):51. PMID: 32272952. doi: 10.1186/s12985-020-01308-y.
173. Constam DB. Regulation of TGFbeta and related signals by precursor processing. Semin Cell Dev Biol. 2014 Aug;32:85–97. PMID: 24508081. doi: 10.1016/j.semcdb.2014.01.008.
174. Ouyang X, Shi X, Huang N, Yang Y, Zhao W, Guo W, et al. WDR72 Enhances the Stemness of Lung Cancer Cells by Activating the AKT/HIF-1alpha Signaling Pathway. J Oncol. 2022;2022:5059588. PMID: 36385964. doi: 10.1155/2022/5059588.
175. LeBlanc ME, Wang W, Caberoy NB, Chen X, Guo F, Alvarado G, et al. Hepatoma-derived growth factor-related protein-3 is a novel angiogenic factor. PLoS One. 2015;10(5):e0127904. PMID: 25996149. doi: 10.1371/journal.pone.0127904.
176. Richter K, Wirta V, Dahl L, Bruce S, Lundeberg J, Carlsson L, et al. Global gene expression analyses of hematopoietic stem cell-like cell lines with inducible Lhx2 expression. BMC Genomics. 2006 Apr 6;7:75. PMID: 16600034. doi: 10.1186/1471-2164-7-75.
177. Qu L, Tang Y, Wu J, Yun X, Lo HH, Song L, et al. FBXL16: a new regulator of neuroinflammation and cognition in Alzheimer's disease through the ubiquitination-dependent

degradation of amyloid precursor protein. Biomark Res. 2024 Nov 21;12(1):144. PMID: 39568047. doi: 10.1186/s40364-024-00691-w.
178. Dou Z, Son JE, Hui CC. Irx3 and Irx5 - Novel Regulatory Factors of Postnatal Hypothalamic Neurogenesis. Front Neurosci. 2021;15:763856. PMID: 34795556. doi: 10.3389/fnins.2021.763856.
179. Fedorova MS, Snezhkina AV, Lipatova AV, Pavlov VS, Kobelyatskaya AA, Guvatova ZG, et al. NETO2 Is Deregulated in Breast, Prostate, and Colorectal Cancer and Participates in Cellular Signaling. Front Genet. 2020;11:594933. PMID: 33362854. doi: 10.3389/fgene.2020.594933.
180. Sahu SK, Fritz A, Tiwari N, Kovacs Z, Pouya A, Wullner V, et al. TOX3 regulates neural progenitor identity. Biochim Biophys Acta. 2016 Jul;1859(7):833–40. PMID: 27080130. doi: 10.1016/j.bbagrm.2016.04.005.
181. Quek LS, Grasset N, Jasmen JB, Robinson KS, Bellanger S. Dual Role of the Anaphase Promoting Complex/Cyclosome in Regulating Stemness and Differentiation in Human Primary Keratinocytes. J Invest Dermatol. 2018 Aug;138(8):1851–61. PMID: 29526760. doi: 10.1016/j.jid.2018.02.033.
182. Rausch P, Steck N, Suwandi A, Seidel JA, Kunzel S, Bhullar K, et al. Expression of the Blood-Group-Related Gene B4galnt2 Alters Susceptibility to Salmonella Infection. PLoS Pathog. 2015 Jul;11(7):e1005008. PMID: 26133982. doi: 10.1371/journal.ppat.1005008.
183. Zhao M, Nakada Y, Wei Y, Bian W, Chu Y, Borovjagin AV, et al. Cyclin D2 Overexpression Enhances the Efficacy of Human Induced Pluripotent Stem Cell-Derived Cardiomyocytes for Myocardial Repair in a Swine Model of Myocardial Infarction. Circulation. 2021 Jul 20;144(3):210–28. PMID: 33951921. doi: 10.1161/CIRCULATIONAHA.120.049497.
184. Xiao Y, Yang Y, Xiong H, Dong G. The implications of FASN in immune cell biology and related diseases. Cell death & disease. 2024 Jan 25;15(1):88. PMID: 38272906. doi: 10.1038/s41419-024-06463-6.
185. Knobloch M, Braun SM, Zurkirchen L, von Schoultz C, Zamboni N, Arauzo-Bravo MJ, et al. Metabolic control of adult neural stem cell activity by Fasn-dependent lipogenesis. Nature. 2013 Jan 10;493(7431):226–30. PMID: 23201681. doi: 10.1038/nature11689.
186. Zhang YW, Mess J, Aizarani N, Mishra P, Johnson C, Romero-Mulero MC, et al. Hyaluronic acid-GPRC5C signalling promotes dormancy in haematopoietic stem cells. Nat Cell Biol. 2022 Jul;24(7):1038–48. PMID: 35725769. doi: 10.1038/s41556-022-00931-x.
187. Lawson H, Sepulveda C, van de Lagemaat LN, Durko J, Barile M, Tavosanis A, et al. JMJD6 promotes self-renewal and regenerative capacity of hematopoietic stem cells. Blood Adv. 2021 Feb 9;5(3):889–99. PMID: 33560400. doi: 10.1182/bloodadvances.2020002702.
188. Saha SK, Choi HY, Kim BW, Dayem AA, Yang GM, Kim KS, et al. KRT19 directly interacts with beta-catenin/RAC1 complex to regulate NUMB-dependent NOTCH signaling pathway and breast cancer properties. Oncogene. 2017 Jan 19;36(3):332–49. PMID: 27345400. doi: 10.1038/onc.2016.221.
189. Saha SK, Yin Y, Chae HS, Cho SG. Opposing Regulation of Cancer Properties via KRT19-Mediated Differential Modulation of Wnt/beta-Catenin/Notch Signaling in Breast and Colon Cancers. Cancers (Basel). 2019 Jan 15;11(1). PMID: 30650643. doi: 10.3390/cancers11010099.
190. Kyrousi C, O'Neill AC, Brazovskaja A, He Z, Kielkowski P, Coquand L, et al. Extracellular LGALS3BP regulates neural progenitor position and relates to human cortical complexity. Nat Commun. 2021 Nov 2;12(1):6298. PMID: 34728600. doi: 10.1038/s41467-021-26447-w.
191. Stampolidis P, Ullrich A, Iacobelli S. LGALS3BP, lectin galactoside-binding soluble 3 binding protein, promotes oncogenic cellular events impeded by antibody intervention. Oncogene. 2015 Jan 2;34(1):39–52. PMID: 24362527. doi: 10.1038/onc.2013.548.
